# Medial septum neurokinin- and somatostatin-sensitive mechanisms mediate sensorimotor and nociceptive behaviours

**DOI:** 10.1101/2020.10.27.358267

**Authors:** Si Yun Ng, Andy Thiam-Huat Lee, Mohammed Zacky Ariffin, Pei Jun Woon, Terence Sheng Hao Chng, Sanjay Khanna

## Abstract

The forebrain medial septum (MS), implicated in affective-motivational behaviours, is enriched in substance P (SP) sensitive neurokinin-1 receptors (NK1R) and somatostatin (SST) receptors (SSTR) that are located almost exclusively on cholinergic and GABAergic neurons, respectively. However, the physiological function of these receptors is poorly understood. This study characterized the actions of intraseptal SP on electrophysiological indices of septo-hippocampal activation, then utilised NK1 receptor antagonist, L-733,060, and SST to investigate the physiological role of endogenous neurotransmission at NK1R, and SST-sensitive mechanisms, in novel open field and formalin test of inflammatory pain. The findings showed that neurotransmission at NK1R mediates formalin-induced electrophysiological responses in the septo-hippocampus in anaesthetized and behaving animals. Furthermore, parallel NK1R- and SST-sensitive mechanisms affect different aspects of animal behaviours in both tests, collectively modulating attention and habituation in open field and driving formalin-induced nociception. This brings out a newer peptidergic dimension of septal physiology in nociception.

## Introduction

The cholinergic neurons in the basal forebrain region of medial septum (MS) modulate a variety of adaptive behaviours, including learning and memory, extinction, latent inhibition, peripheral injury-induced hyperalgesia and allodynia, response to anaesthetics and anxiety (Baxter *et al*., 1997; Easton *et al*., 2011; Jiang *et al*., 2018b; Marston *et al*., 1994; Tronson *et al*., 2009; Tai et al., 2014; Zhang *et al*., 2017). Consistently, different lines of evidences indicate that septal cholinergic neurons are excited to behaviourally salient stimuli. For example, neural activity of septal cholinergic neurons is increased on peripheral sensory stimulation or on exposure to novel environment (Ceccarelli *et al*., 1999; Giovannini *et al*., 2001; Lovett-Barron *et al*., 2014). In turn, these neurons promote signalling in downstream structures, especially hippocampus, to salient stimuli including hind paw injection of the algogen formalin, a model of persistent nociception (Brazhnik *et al*., 2004; Khanna, 1997; Lovett-Barron *et al*., 2014; Zheng & Khanna, 2001). The septal cholinergic neurons also facilitate hippocampal theta activation, a 3-12 Hz sinusoidal extracellular activity, reflecting a theta-rhythmic synchronized modulation of hippocampal neurons (Bland, 1986; Buzsáki, 2002; Dannenberg *et al*., 2015; Gasparini & Magee, 2006; Lee *et al*., 1994; Leung, 1984; Magee, 2001; Mamad *et al*., 2015; Tai *et al*., 2006; Vandecasteele *et al*., 2014; Zheng & Khanna, 2001).

The septal cholinergic neurons are enriched in neurokinin receptors (NKR) including, NK1R NK2R and NK3R, which modulate their excitability (Chen *et al*., 2001; de Souza Silva *et al*., 2013; Morozova *et al*., 2008; Schäble *et al*., 2012; Steinberg *et al*., 1998). Indeed, the NKR are almost exclusively localized to septal cholinergic neurons. Further, post-conditioning administration of SP into MS in behaving animals facilitates learning of avoidance behaviour (Siegel *et al*., 1984; Stäubli & Huston, 1980). However, while the septal cholinergic neurons, and more generally the cholinergic neurons of the basal forebrain, facilitate attention/arousal (Fuller *et al*., 2011; Tsanov, 2017; Vandecasteele *et al*., 2014; Zant *et al*., 2016), the role of NKR in modulation of acute behaviour to salient stimuli remains undefined.

In a striking contrast to NKR, the peptidergic somatostatin (SST) receptors (SST_2A_R) are expressed on a proportion of GABAergic, but not cholinergic neurons of the MS, suggesting that SST exercises a differential effect on septal circuits vis-à-vis transmission at NK1R (Bassant *et al*., 2005). Arising from the preceding, the present study has examined the influence of septal peptidergic transmission at NK1R vis-à-vis SST-sensitive mechanisms on animal behaviour and hippocampal theta activation in novel environment and the formalin test. Hippocampus is implicated in pain. Indeed, formalin induces septal cholinergic neuron-mediated hippocampal theta activation during which the excitation of CA1 pyramidal cells is sculpted in a theta rhythmic fashion (Tai *et al*., 2006; Zheng & Khanna, 2001). Further, both animal models display behavioural activation that varies with time allowing for an exploration of effect at different phases of behaviour.

## Materials and Methods

### Animals

The experimental studies described here were approved by the local institutional Animal Care and Use Committee (IACUC) from National University of Singapore (NUS). Experiments were performed on adult male Sprague Dawley rats aged 7-8 weeks old (270 to 350 g) purchased from Laboratory Animals Center and InVivos Pte Ltd. Rats were housed under controlled temperature and humidity in the animal facility with water *ad libitum* and pelleted food under a 12hr dark/light cycle.

### Surgical procedures

#### Anaesthetized animals

Urethane (1 g/kg, i.p.; Sigma, USA) anaesthetized animals were mounted onto a stereotaxic frame (Stoelting Co, Wood Dale, IL, USA). A burr hole was made overlying the MS (A0.5 mm from Bregma, L0.5 mm from midline, and V6.5 mm from the cortical surface) (Paxinos & Watson, 2007) to facilitate lowering of a microsyringe or a double-barreled cannula. A bone flap was made over the left cerebral hemisphere for lowering of recording and stimulating electrodes (Lee *et al*., 2011).

#### Cortical implantations

Survival surgery was performed under aseptic conditions using stereotaxic technique as previously described (Ang *et al*., 2015; Ariffin *et al*., 2018; Lee *et al*., 2011). Briefly, anesthesia was induced and maintained with 5% and 2% isoflurane, respectively. Oxygen was provided at 1 litre/min. A single barreled 26G stainless steel guide cannula (Plastic One, Roanoke, VA, USA) was directed towards the dorsal edge of MS (A0.5 mm from Bregma, 0.0 mm from midline and V5.8 mm from the cortical surface; Paxinos and Watson, 2007). Animals were also implanted with a depth recording electrode, constructed by twisting a pair of stainless steel Teflon-insulated wires (A-M Systems, USA; 125 μm in diameter) with gold plated male connector pins. The recording electrode was directed towards the stratum radiatum of the left hippocampal field CA1 (P3.0 mm to Bregma, L2.4 mm from midline, and V3.0 mm from the cortical surface; Paxinos and Watson, 2007). Implants were lowered through the cortex via burr holes made on the skull and secured with support screws and dental cement. Following surgical procedure, the animals were housed individually until further experimentation.

Post-operative, the animals were allowed to recover for at least 7 days, during which they were treated with the analgesic buprenorphine (0.03 mg/kg for 3 days, i.p.) and the antibiotic enrofloxacin (10 mg/kg for 5 days, i.p.).

### *In vivo* intracerebral stimulation and electrophysiological recording

#### Anaesthetized animals

A concentric bipolar stimulating electrode (Model NE-100, David Kopf, USA) was directed towards the left hippocampal field CA3 (P3.0 mm from Bregma, L2.4 mm from midline, and V4.0 mm from the cortical surface; Paxinos and Watson, 2007). The CA3 region was stimulated (0.1Hz, 0.01s pulse duration) through a constant current stimulation isolation unit (Grass S88 stimulator, Grass technologies, Warwick, RI, USA) to evoke a population spike (PS) in the pyramidal layer of the ipsilateral CA1 region. Stimulation intensity was adjusted to generate a PS amplitude at 70% of the maximum (Ariffin *et al*., 2010; Lee *et al*., 2011).

The PS was recorded in parallel with local extracellular field activity via a saline filled carbon fiber glass microelectrode (Jiang & Khanna, 2004, 2006; Lee *et al*., 2011; Zheng & Khanna, 2001). The recording electrode was oriented at an angle of 5° to right from the vertical and directed towards the pyramidal layer of the left hippocampal field CA1 (P3.6 mm from Bregma, L2.0 mm from midline, and V4.0 mm from the cortical surface; Paxinos and Watson, 2007). The electrode position in or around pyramidal cell layer was identified by the observation of complex spike cells and PS. As previously (Khanna, 1997), the signal from the carbon fiber was amplified (Grass amplifier, Astromed Inc., USA) and differentially filtered at (a) 1–100 Hz, to record local hippocampal field activity, and (b) 1–3000 Hz to record PS. Data were digitized (Power 1401 A/D convertor, Cambridge Electronic Design, UK) at 10 KHz for PS and 256 Hz for field activity and collected using Spike 2 software (Cambridge Electronic Design, UK) for offline analysis.

#### Awake animals

To record hippocampal field activity in behaving animals, a flexible recording wire was connected to the implanted head stage on the animal while the other end was connected to a commutator that was suspended above the test chamber. The field activity signals were amplified 500x and digitized at 256 Hz. Signals were band pass filtered between 1-100 Hz, digitized as above and recorded with the Spike 2 program.

### Drugs and microinjection procedure

The drugs used were i) the NK1R agonist, Substance P (SP; 1 and 2 μg/μl; #S6883, Sigma, USA), ii) the NK1R antagonist, L-733,060 (1X: 0.0176 μg/μl and 10X: 0.176 μg/μl; #1145, Tocris Bioscience, UK), iii) the cholinergic receptor agonist, carbamoylcholine chloride (carbachol; 0.156 μg/μl; #C4382, Sigma, USA) and iv) the peptide neurotransmitter somatostatin (SST; 3.38 μg/μl and 6.76 μg/μl; Bachem, USA). All drugs were dissolved in saline (0.9% w/v sodium chloride; Sigma, USA) in Alcian blue dye (0.5% w/v; Sigma, USA). The dye solution was administered as the corresponding vehicle. All drugs were microinjected into the septum in a volume of 0.5 μl.

The functional effects of different doses of SP and L-733,060 were explored in this study. Notably, L-733,060 at a concentration explored in the present study is anti-nociceptive on microinjection into the rostral ventral medulla (Hamity *et al*., 2010). The concentration of carbachol used in the experiments is based on published work (Jiang & Khanna, 2006). The concentration of SST was selected from published work wherein intraseptal microinjection of the peptide at the selected concentrations disrupted theta wave activity during exploration in an open field (Bassant *et al*., 2005).

#### Anaesthetized animals

The drugs were administered into the MS either via a 33G stainless steel microinjection needle attached to an Exmire microsyringe (Ito Corporation, Fuji, Japan) or via a double-barrelled cannula system. The microinjection needle/cannula was lowered into the MS at an angle of 5° to the right from the vertical. The double barrel was orientated along the anterior-posterior axis of the MS. The antagonist, L-733,060, or vehicle was microinjected via the anterior cannula while SP or carbachol was microinjected via the posterior cannula.

#### Awake animals

Drugs were administered via a 33G stainless steel internal cannula (Plastic One, Roanoke, VA, USA) which was connected to a 25 μl microsyringe (Hamilton, USA) by a polyethylene cannula connector assembly system. The internal cannula protrudes from the tip of the guide cannula by 1.0 mm. With gentle restraint, the internal cannula was inserted into the implanted guide cannula and the drug was microinjected over a period of 30s. The internal cannula was left in situ for at least 1 minute to allow diffusion and to minimize backflow. Drugs were administered in a blinded fashion.

### Histology

At the end of each experiment, the animals were given an overdose of urethane (1.5 g/kg, i.p., Sigma, USA). The animals were perfused transcardially with 0.9% sodium chloride solution followed by 10% formalin (Merck, Germany). The brain was removed and placed in the fixative for later sectioning. Tissues were sectioned on a vibratome (Leica VT1200, Leica Microsystems GmbH, Wetzlar, Germany) and collected in Tris buffered saline (TBS). Alternate 100 μm coronal sections were collected for Nissl-staining (0.5% w/v Cresyl violet, Sigma, USA) to determine stimulating, recording and microinjection sites.

### Experimental protocol

#### Investigation into the selectivity of antagonism by NK1R antagonist L-733,060

Different concentrations of L-733,060 (0.5 μl of 0.0176 μg/μl [1x] or 0.176 μg/μl [10x]) were microinjected into MS in separate experiments to investigate the ability of the drug to antagonize hippocampal responses evoked on intraseptal microinjection of either SP or carbachol. SP is an agonist at NK1R, while carbachol is an agonist at cholinergic muscarinic receptor. A double-barrelled cannula was used in these experiments. The concentration of SP (0.5 μl of 2 μg/μl) used in these experiments was determined in a pilot study. In that study, different concentrations of SP were used to find a low concentration that induced robust and reproducible effect on amplitude of hippocampal CA1 PS, while the concentration of carbachol used in the experiments is based on published work (0.5 μl of 0.156 μg/μl; Jiang and Khanna, 2006). Microinjection of carbachol into MS has been shown to elicit robust suppression of CA1 PS and evoke a strong theta activation in hippocampal CA1 (Zheng & Khanna, 2001).

Each experiment comprised of three microinjections of the agonist, at least 60 min apart. L-733,060 or vehicle was microinjected only once, 15 min before the second microinjection of the agonist, under condition of large irregular hippocampal field activity (LIA). Because the second microinjection was preceded by microinjection of the antagonist or vehicle, the data was labelled as either L-733,060 + SP2, L-733,060 + C2, Vehicle + SP2 or Vehicle + C2 where L-733,060 and Vehicle indicate the antagonist and vehicle, respectively, while SP2 and C2 signify the 2^nd^ microinjection of SP and carbachol, respectively. Further, a prefix such as 1x was added before L-733,060 to identify the concentration of the antagonist used in the experiment. The responses to the first and third microinjection of SP and carbachol were labelled as SP1 and SP3, and C1 and C3, respectively.

The effects of agonists were monitored continually for 20 min after microinjection. Additional 2 min recordings were made at 20 min interval up to 60 min post microinjection.

#### Effect of intraseptal L-733,060 on formalin-induced nociception

The characterisation of effect of NK1R on formalin-induced electrophysiological responses was first investigated in anaesthetized animal. Based on the previous experiment (see above), the concentration of L-733,060 selected for the study was 0.0176 μg/μl (1x; 0.5 μl). L-733,060 was microinjected into septum 15 min preceding intra-plantar injection of formalin (5%, 0.05 ml) into right hind paw. The electrophysiological responses were continually monitored at time points before microinjection, following microinjection and post-formalin injection. Post-formalin injection, the hippocampal responses were recorded continuously for 20 minutes and for 2 minutes at the 40^th^ and 60^th^ minute.

#### Effect of intraseptal L-733,060 on exploratory behaviors in the open field

In these experiments, the effect of intraseptal microinjection of a drug (0.5 μl of either 0.0176 μg/μl of L-733,060, 3.38 μg/μl or 6.76 μg/μl of SST, or vehicle) was examined on animal ambulation and hippocampal theta field activity recorded concurrently in the novel open field. The open field behavior was recorded after 15min and immediately after microinjection of L-733,060 and SST, respectively. As previously (Ang *et al*., 2015), the animals were placed at the lower left corner of the open field arena (L43.2 cm × W43.2 cm × H30.5 cm, L×W×H; Model ENV-515, Med Associates Inc., USA) located in a behavioral suite. The behavior was monitored for 60 min. Animal’s movements and position were tracked by the open field monitor using 3 sets of 16 evenly spaced infrared (IR) transmitters and receivers positioned along the perimeter of the chamber. Ambulatory distance and speed were computed by the Actimot software (Med Associates Inc, USA). An ambulatory episode was signaled when the distance travelled by the animal exceeded the pre-set space of 4 x 4 IR beams (chamber size of approximately 10 cm by 10 cm). The speed was measured by the software if the animal moved for more than 2s.

Control theta wave activity was recorded for at least 2 min during animal exploration in a familiar laboratory adjacent to the behavioral suite. The power of control theta was used to normalize the power of theta induced during exploration of novel environment.

#### Effect of intraseptal L-733,060 or SST on formalin-induced nociceptive responses

The formalin test was conducted in a test chamber (L43.2 cm x W21.7 cm x H30.5 cm) of an open field activity monitor (model ENV-515, Med Associates Inc., see above). Prior to the day of the formalin test, animals were habituated to the experimental chamber for at least 60 min each day for 3 consecutive days. On the day of formalin test, animals were generally habituated for 30 min before drug microinjection and hind paw injection of formalin.

Prior to the experiment, control hippocampal theta wave activity was recorded for at least 2 min during exploration of the test arena by the animal. The power of theta was used to normalize the power of theta induced following injection of formalin. The exploration of the test arena was followed by microinjection of 0.5 μl of either L-733,060 (0.0176 μg/μl or 0.176 μg/μl), SST (3.38 μg/μl or 6.76 μg/μl, 0.5 μl) or vehicle. The formalin test was either performed immediately after microinjection of SST or the corresponding vehicle, or 15 min after microinjection of L733,060 or the associated vehicle. This 15 min period before formalin injection is labelled as the baseline recording during which the animal was allowed to spontaneously explore the test chamber. The formalin test involved injecting formalin (1.25%, 0.1 ml) subcutaneously into the plantar surface of the right hind paw.

Formalin-induced nociceptive behaviours were monitored by the experimenter for a period of 60 min. Nociceptive licking was measured as the time spent licking the injured paw at both the dorsal and plantar surfaces. While, nociceptive flinching was the number of shakes and jerks of the injured hind paw. In addition, formalin-induced ambulation (moment-to-moment agitation) was also measured by recording animal ambulatory movements using the open field activity monitor (see above). The hippocampal field activity was recorded using Spike 2 software.

### Data Analyses

#### Electrophysiological recording

PS and hippocampal theta wave activity were analyzed offline using Spike 2 software. PS amplitude (mV) was calculated as the average of the negative peak from the 2 positive peaks around it as previously described (Ariffin *et al*., 2010; Jiang & Khanna, 2004, 2006; Khanna, 1997; Lee *et al*., 2011). PS amplitude was averaged over six sweeps in 1 min blocks. The magnitude of PS amplitude reflects the size of the neuronal population discharging synchronously in response to CA3 stimulation.

The hippocampal theta wave activity was analyzed for the following parameters: i) theta duration (s/min), ii) peak theta frequency (Hz), and iii) normalized peak theta power. The peak theta frequency and power refers to the fast Fourier transform (FFT) theta peak frequency and FFT theta peak power in the theta frequency range of 3-6 Hz in anaesthetized animals or 4-12 Hz in behaving animals. Field activity was analyzed in blocks of 1 min and 5 min in anaesthetized and behaving animals, respectively.

During analysis, the extracellular field activity trace was digitally band-pass filtered at 1-40 Hz with finite impulse response (FIR) filter. Theta segments of at least 2 s duration from the FIR filtered data were used for FFT analyses (frequency resolution of 0.5 Hz) to derive the average FFT theta peak frequency (Hz) and FFT theta peak power in 1 min blocks. As described previously (Tai *et al*., 2006), the FFT theta power presented as units of mV^2^ by the program was normalized to give the power (peak-to-peak amplitude square in the mV^2^ unit) of the field activity. The final computed average FFT theta peak power was normalized to the average FFT theta peak power derived from the baseline recording in each experiment. In the anaesthetized experiments, theta power was normalized to the power of spontaneous theta activity recorded during baseline, while in the behaving animals, theta power was normalized with the theta power computed from exploration-induced theta recorded prior to the experiment. Duration of theta was calculated by computing the period of time (s per 1 or 5 min block) for which theta was visually identified as a continuous sinusoidal oscillation of at least 1s duration at frequencies of 3–12 Hz.

#### Open field

The total ambulatory distance (cm) and average speed (cm/s) were extracted from the Actimot software in 5 min blocks.

#### Formalin-induced responses

The total ambulatory distance (cm) in 5 min blocks was extracted from the Actimot software. Formalin-induced flinches (counts/5min) and duration of licking (s/5min) of the injured paw were summed in 5 min blocks. Phase analysis of formalin-induced responses was carried out.

In this regard, phase 1 refers to the 1-5 min period and phase 2 refers to the 11-60 min period, while interphase represents 6-10 min period.

### Statistical analysis

Based on previous studies from the laboratory, we anticipated that typically 6 to 12 animals in a group are required for behavioural and electrophysiological experiments to pinpoint clear differences in data, if any, among groups (Ang *et al*., 2015; Lee *et al*., 2011). Statistical analysis of data was carried out using Prism 4 (GraphPad Software, USA). The time course graphs depicting the changes in the electrophysiological or behavioural parameters to a treatment were analysed using one-way repeated measure (RM) ANOVA followed by Newman-Keuls post-hoc test. Whereas, the difference in the time course of change between treatments was compared with two-way RM ANOVA followed by Bonferroni post-hoc test. The difference between means of multiple groups were analysed using one-way ANOVA followed by Newman-Keuls post-hoc test. In instances when the Bartlett’s test showed unequal variance, the data was normalized by logarithmic transformation. An ANOVA was used for analysis if the transformed data exhibited equal variance. Otherwise, the non-parametric Kruskal-Wallis test was applied to the data. Differences were considered to be statistically significant at p ≤ 0.05. The data are expressed and represented as mean ± standard error of mean (S.E.M).

## Results

### Pharmacological investigation with drug microinjection into medial septum in anaesthetized rat

The NK1R antagonist, L-733,060, was pre-administered 15min prior to other pharmacological and physiological manipulations. The drug was administered as pre-treatment so as to determine its effect, if any, on the basal electrophysiological responses.

### Population spike

#### Effect of microinjection of the NK1R antagonist, L-733,060, on suppression of hippocampal CA1 population spike (PS) induced by intraseptal SP

<u>Dose dependent effect of SP</u>: The dose of SP (2 μg/μl) was selected based on a pilot study in which different doses of SP (1 μg/μl, n = 7; 2 μg/μl, n = 9) or vehicle (n = 5) was microinjected into MS in separate experiments (Figure S1A). In another experiment, SP at 2 μg/μl was microinjected into LS (n = 7). SP was microinjected thrice, at least 1 hr apart. Since the effect of repeat injections at a given dose and at the selected site was comparable, an average response for that dose and site was built by averaging the time course for the three microinjections in a given experiment and then for the entire group. Compared to vehicle treatment, PS was suppressed only on microinjection of SP into MS, but not LS (Figure S1B; Treatment, F_3, 600_ = 16.97, p < 0.0001; two-way RM ANOVA followed by Bonferroni post-hoc test; also see below). Comparison of the average amplitude of PS in the first five minutes after drug microinjection, expressed as percentage of the control amplitude before microinjection, indicated that the suppression evoked on microinjection of the higher dose of SP into MS was more robust as compared to the lower dose (Figure S1B; Groups, p < 0.0001; Kruskal-Wallis test followed by Dunn’s post-hoc test). The control amplitude, i.e. average amplitude of the control PS in the two minutes (i.e. from −2-0 min on left plot, Figure S1B) preceding microinjection of vehicle or the SP was similar between the groups (Groups, F_3, 24_ = 1.86, p > 0.1; one-way ANOVA). The control amplitudes for Vehicle vs. SP (2 μg/μl) in LS vs. SP (1 μg/μl) in MS vs. SP (2 μg/μl) in MS were 6.25 ± 0.13 mV (n=5) vs. 6.30 ± 0.30 mV (n=7) vs. 6.93 ± 0.25 mV (n=7) vs. 6.48 ± 0.16 mV (n=9).

#### Antagonistic effect of L-733,060 on SP-induced suppression

SP was microinjected thrice, at least 1 hr apart, in the study to investigate the sensitivity of SP-evoked responses to antagonism by the NK1R antagonist, L-733,060. The antagonist (1x = 0.00176 μg/μl, or 10x = 0.0176 μg/μl) or the corresponding vehicle was microinjected 15min before the 2^nd^ microinjection of SP (Figure 1A, B).

**Figure 1.**
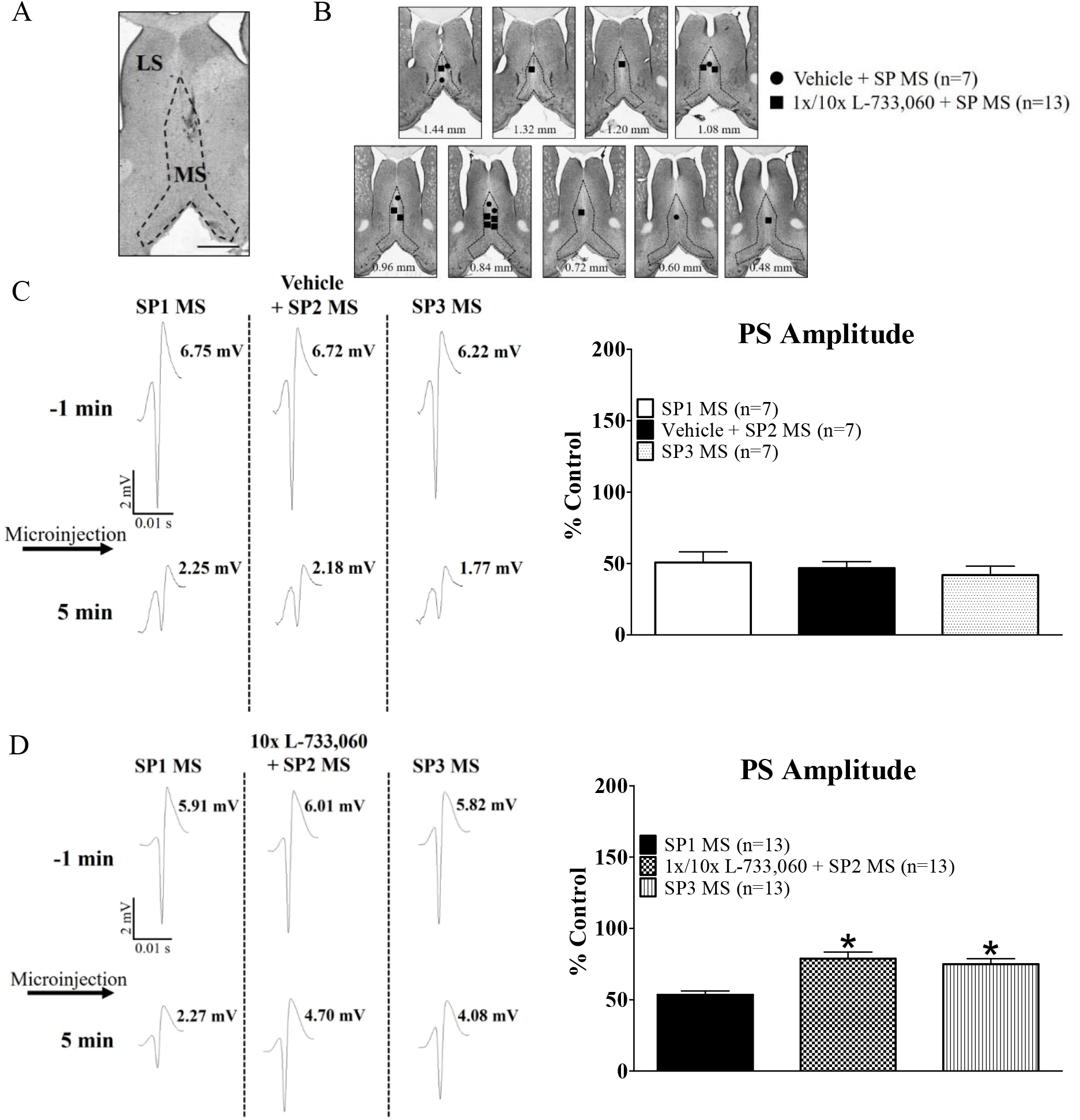
Microinjection of neurokinin 1 receptor (NK1R) antagonist, L-733,060, into the medial septum (MS) blocked Substance P (SP)-induced suppression of CA1 population spike (PS) in anaesthetised animals. (A) Representative microinjection site in the MS. (B) Composite of microinjection sites in the MS. SP (0.5 μl of 2 μg/μl) was microinjected via a double-barrelled silicon cannula with vehicle or L-733,060 (1x, 0.0176 μg/μl; 10x, 0.176 μg/μl) in separate experiments. (C) The NK1R agonist SP was microinjected three times (SP1, SP2, SP3), whereas vehicle or L-733,060 was microinjected 15 min before SP2. The PS traces on the left represent the average PS in 1-min block before (−1 min) and after (5 min) SP microinjection. The histogram on the right is the average CA1 PS amplitude in 5 min after SP microinjection expressed as percentage of control amplitude before microinjection. (C) Notice the robust and consistent decrease in PS amplitude on SP microinjection, with no effect of vehicle pre-treatment. (D) Pre-treatment with L-733,060 attenuated SP-induced suppression of CA1 PS. Data are presented are mean + S.E.M. Significant difference (p < 0.05): * vs. SP1 MS; one-way ANOVA followed by Newman-Keuls post-hoc test.

The three repeat microinjections of SP were labelled as: SP1 for 1^st^ microinjection; vehicle or different doses of L-733,060 + SP2 for 2^nd^ microinjection; and SP3 for 3^rd^ microinjection. Because of pharmacological intervention with the antagonist, the time courses of effect with repeat microinjections of SP were not averaged unlike that in the pilot study.

The ‘control’ effect with repeated microinjections of SP in vehicle pre-treated group was marked by robust suppression of CA1 PS vis-à-vis the amplitude in the two minute prior to the microinjection (SP1: Time, F_23, 138_ = 8.55, p < 0.0001; vehicle + SP2: Time, F_23, 138_ = 14.63, p < 0.0001; SP3: Time, F_23, 138_ = 9.88, p < 0.0001; n = 7, one-way RM ANOVA followed by Newman-Keuls post-hoc test; data not shown). The suppression was significant for at least 30 min after microinjection of SP.

A two-way RM ANOVA of the time points post microinjection of SP showed that the amplitudes of PS were similar indicating that a similar pattern of time course of suppression is observed with repeated microinjections of SP in vehicle pre-treatment group (Figure S2A; Treatment, F_2, 414_ = 0.24, p > 0.7). Indeed, the time course plots overlapped.

The average percentage of PS amplitude was also very similar across the repeated microinjections (Figure 1C; Groups, F_2, 18_ = 0.48, p > 0.6; one-way ANOVA). The control amplitude, i.e. average amplitude of the control PS in the two minutes preceding microinjection of SP was also similar (Groups, F_2, 18_ = 1.18, p > 0.3). The control amplitudes for SP1 vs. vehicle + SP2 vs. SP3 were 6.04 ± 0.20 mV vs. 5.94 ± 0.15 mV vs. 5.67 ± 0.17 mV (n = 7).

In contrast, pre-treatment with different doses of L-733,060 in separate experiments significantly attenuated SP-induced suppression. In the analysis, the time courses of change in a group after repeat microinjections of SP were compared using two-way RM ANOVA followed by Bonferroni post-hoc test. The analysis showed a significant effect of pre-treatment of different doses of L-733,060 (1x L-733,060, Treatment, F_2, 441_ = 5.38, p < 0.02, n = 8; 10x L-733,060, Treatment, F_2, 252_ = 23.62, p < 0.0001, n = 5; data not shown).

Between group comparison of the time courses after microinjection of SP revealed comparable responses in the two groups (1x L733,060 pre-treatment group vs. 10x L733,060 pre-treatment group: SP1 vs. SP1, Treatment, F_1, 231_ = 4.66, p > 0.05; 1x L733,060 + SP2 vs. 10x L733,060 + SP2, Treatment, F_1, 231_ = 0.23, p > 0.6; SP3 vs. SP3, Treatment, F_1, 231_ =0.87, p > 0.3; twoway RM ANOVA; data not shown). As a result, the two groups were combined.

A comparison of the post SP microinjection time points in the combined group revealed an attenuation of PS suppression at SP2 and SP3 compared to SP1 (Figure S2A; Treatment, F_2, 756_ = 15.60, p < 0.02; two-way RM ANOVA followed by Bonferroni post-hoc test). The attenuation was significant from 2^nd^ to 20^th^ min at SP2, and at 2^nd^ min and at time points from 11^th^ to 20^th^ min at SP3. Generally, the suppression with SP per se is significant for about 30min (see above under the effect on PS in vehicle pre-treated group).

Consistently, a comparison showed that the average percentage of PS amplitudes with SP2 and SP3 was significantly higher than that observed with SP1 (Figure 1D; Groups. F_2, 36_ = 12.93, p < 0.0001; one-way ANOVA followed by Newman-Keuls post-hoc test). This suggests that the suppression evoked by SP was attenuated at SP2 and SP3 following pre-treatment with L-733,060.

#### Lack of effect of L-733,060 on control amplitude

In the L-733,060 pre-treated groups, the average amplitude of the CA1 PS in the two minutes preceding 2^nd^ microinjection of SP (SP2) was no different from the corresponding amplitudes before 1^st^ (SP1) and 3^rd^ (SP3) microinjection of SP. The two minutes preceding SP2 corresponded to the last two minutes of the 15min period following microinjection of L-733,060.

In context of above, the control amplitudes of the PS preceding repeat microinjection of SP for 1x L-733,060 pre-treatment group were - SP1 vs. 1x L-733,060 + SP2 vs. SP3: 6.17 ± 0.19 mV vs. 6.49 ± 0.37 mV vs. 6.34 ± 0.26 mV (Groups, F_2, 21_ = 0.32, p > 0.7, n = 8). Likewise, the values for 10x L-733,060 pre-treatment group were - SP1 vs. 10x L-733,060 + SP2 vs. SP3: 5.68 ± 0.19 mV vs. 5.60 ± 0.20 mV vs. 5.55 ± 0.12 mV (Groups, F_2, 12_ = 0.17, p > 0.8, n = 5). The control amplitudes of PS were also very similar across the two groups of L-733,060 pretreated animals (Groups, F_5, 33_ = 2.29, p > 0.06; one-way ANOVA).

#### Effect of microinjection of L-733,060 on suppression induced by intraseptal carbachol

The ‘control’ effect with repeated microinjections of carbachol in vehicle pre-treated group was marked by robust suppression of CA1 PS vis-à-vis the amplitudes in the two minute prior to the microinjection (Figures S2B, 2A and B; C1: Time, F_23, 92_ = 14.42, p < 0.0001; vehicle + C2: Time, F_23, 92_ = 14.75, p < 0.0001; C3: Time, F_23, 92_ = 11.69, p < 0.0001; n = 5, one-way RM ANOVA followed by Newman-Keuls post-hoc test; data not shown). A two-way RM ANOVA of the time courses after microinjection of carbachol showed that the amplitudes of PS were similar indicating that a similar pattern of time course of suppression is observed with repeated microinjections of carbachol in vehicle pre-treatment group (Figure S2C; Treatment, F_2, 252_ = 1.59, p > 0.2). Indeed, the average percentages of PS amplitude were very similar across the repeated microinjections (Figure 2B; Groups, F_2, 4_ = 1.58, p > 0.2; one-way ANOVA, n = 5). The control amplitude, i.e. average amplitude of the control PS in the two minutes preceding microinjection of carbachol was also similar (Groups, F_2, 12_ = 1.45, p > 0.2). The control amplitudes for C1 vs. vehicle + C2 vs. C3 were 6.31 ± 0.28 mV vs. 6.07 ± 0.26 mV vs. 5.68 ± 0.24 mV (n = 5).

**Figure 2.**
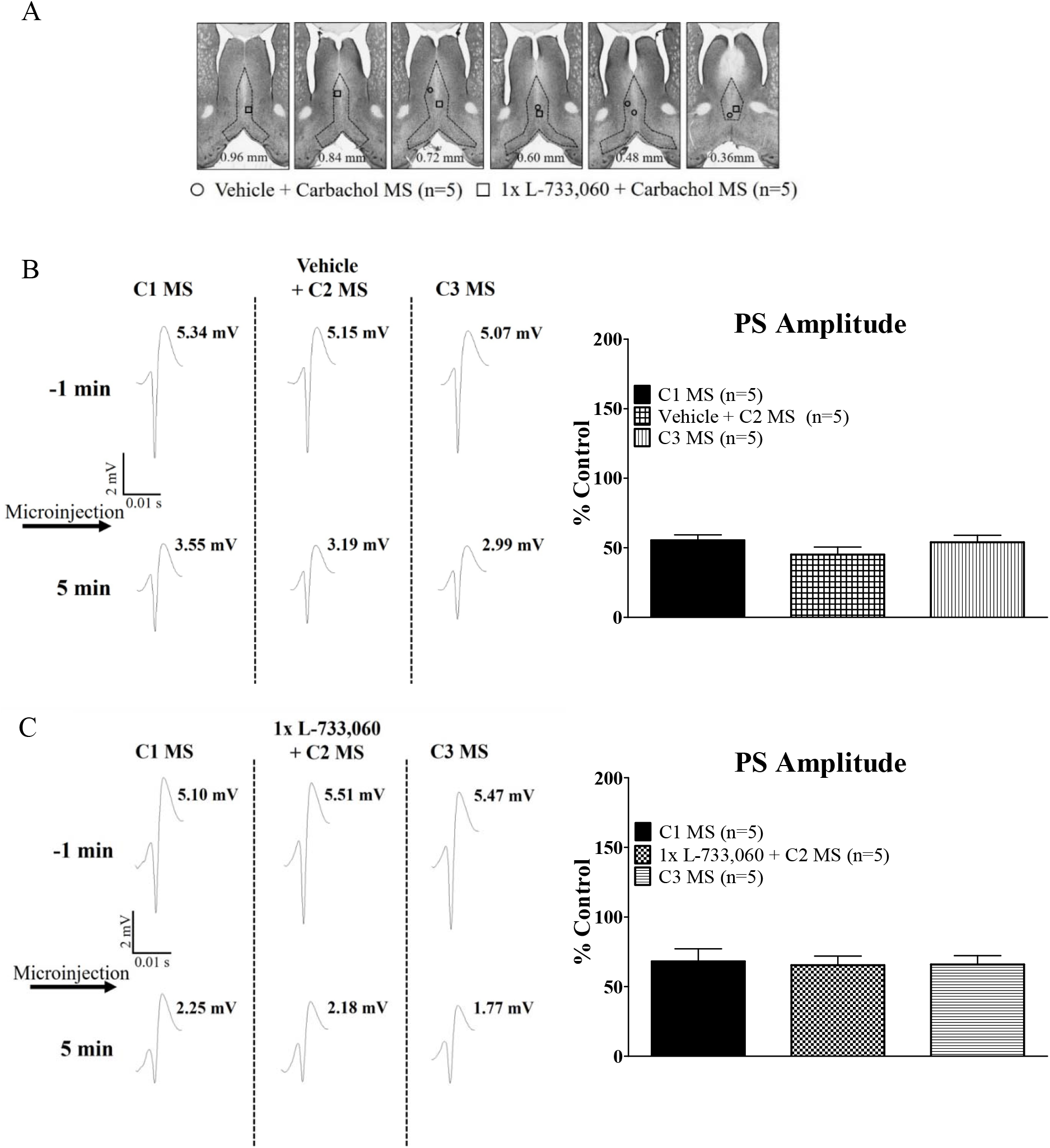
Microinjection of L-733,060 into the medial septum did not affect carbachol-induced suppression of CA1 population spike (PS) in anaesthetized animals. (A) Composite representation of the microinjection sites in the medial septum (MS). The composite is built as explained in Fig. 1. Carbachol (0.0156 μg/μl, 0.5 μl) was microinjected three times (C1, C2 and C3) at least 1hr apart during the course of the experiment whereas vehicle or neurokinin1 receptor (NK1R) antagonist, L-733,060 (1x, 0.0176 μg/μl, 0.5 μl) was microinjected 15 min before the second carbachol administration (C2). The PS traces (left) and histograms (right) are built as explained in Fig. 1. Pre-treatment with vehicle (B) or L-733,060 (C) did not significantly affect PS suppression evoked on carbachol microinjection into MS. Data are presented are mean + S.E.M.

As with vehicle pre-treatment, pre-treatment with 1x L-733,060 did not significantly attenuate carbachol-induced suppression. In the analysis, the time courses of change after repeated microinjections of carbachol were compared using two-way RM ANOVA. The analysis showed an insignificant effect of pre-treatment with 1x L-733,060 (Figure S2B; Treatment, F_2, 252_ = 0.16, p > 0.8). The control amplitudes of the PS preceding repeated microinjection of carbachol were not different from each other (C1 vs. 1x L-733,060 + C2 vs. C3: 5.98 ± 0.48 mV vs. 6.11 ± 0.44 mV vs. 6.06 ± 0.46, n = 5; Groups, F_2, 12_ = 0.02, p > 0.9). The average percentage of PS amplitude was also similar across repeated microinjections of carbachol in this group (Figure 2C; Groups, F_2, 4_ = 0.06, p > 0.9, n = 5; one-way ANOVA).

#### Both agonist induced similar suppression

To discount the possibility that the differences in the effectiveness of L-733,060 against SP vs. carbachol was due to different baselines, the suppression evoked by the two agonists was compared. To this end, the time courses of change with SP1 in the vehicle and the L-733,060 pre-treatment groups were put together (n = 20). Likewise, for C1 (n = 10). Two-way RM ANOVA of the time course of change evoked by S1 vs. C1 revealed an insignificant effect of treatment (Treatment, F_1, 588_ = 0.03, p > 0.8; data not shown), suggesting that PS suppression evoked by the two agonists was similar. The control amplitudes of PS preceding microinjections were also similar (SP1 vs. C1: 6.05 ± 0.13 mV vs. 6.09 ± 0.24 mV, p > 0.8; two-tailed unpaired t-test).

#### Effect of microinjection of L-733,060 on suppression induced by hind paw injection of formalin

To investigate if the formalin-induced suppression is sensitive to antagonism by L-733,060, the algogen was injected 15 min after pre-treatment with microinjection of L-733,060 or vehicle into MS (Figure 3A and B). As previously (Khanna, 1997), injection of formalin evoked a robust and sustained suppression of amplitude of CA1 PS in control experiments involving pre-treatment with vehicle (Time, F_23, 184_ = 6.40, p < 0.0001; one-way RM ANOVA followed by Newman Keuls post-hoc test, n = 9; data not shown). A significant suppression was observed till 60^th^ min after formalin injection.

**Figure 3.**
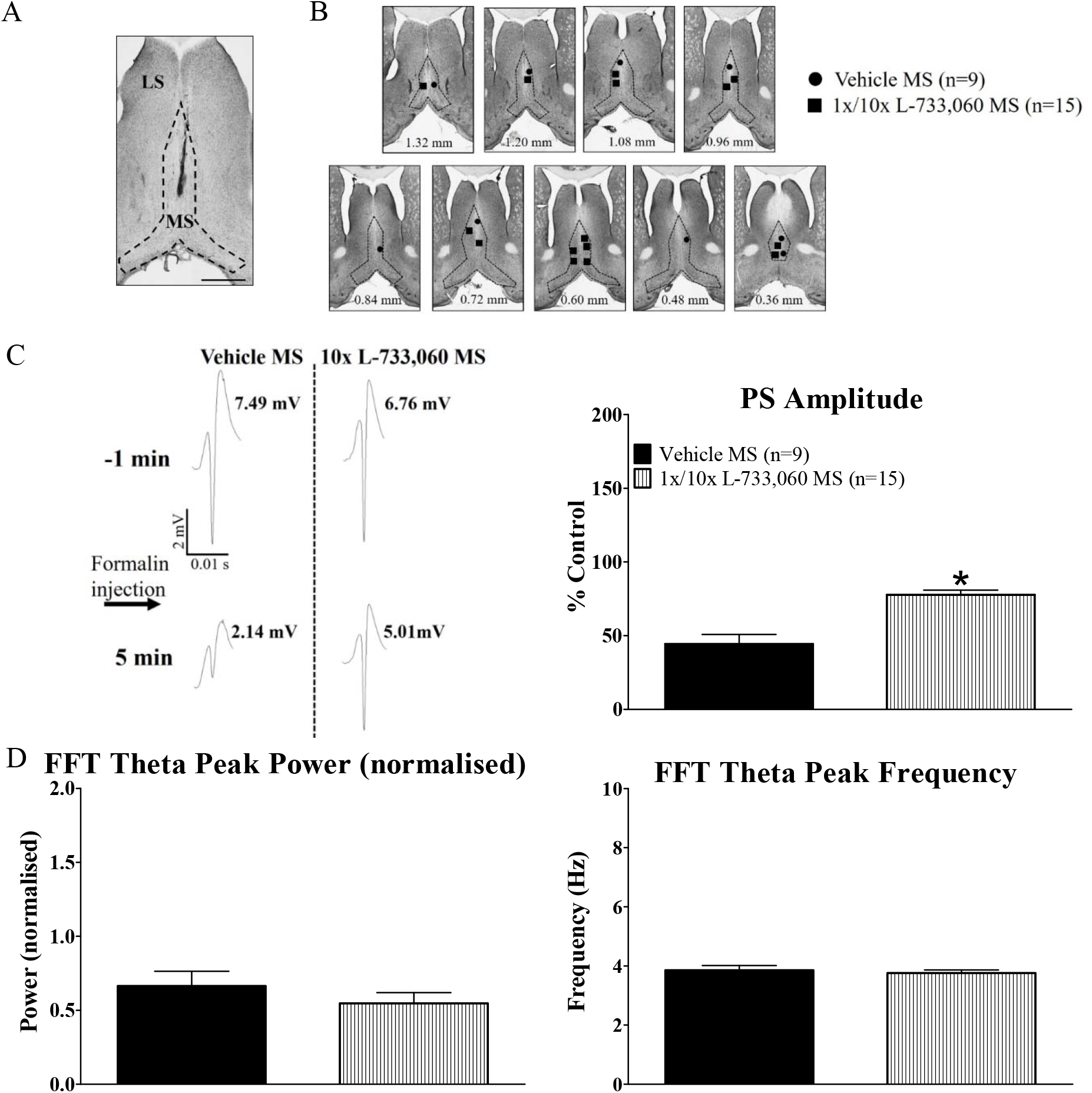
Microinjection of neurokinin 1 receptor (NK1R) antagonist, L-733,060, into the medial septum (MS) attenuated formalin-evoked suppression of CA1 population spike (PS) in anaesthetized animals. (A) Representative microinjection site in the MS. The drug was microinjected into the MS via a single 33G microinjection stainless steel needle coupled to a microsyringe. (B) Composite representation of microinjection sites in MS built as explained in Fig. 1. Formalin (5%, 0.05ml) was injected 15 min after microinjection of NK1R antagonist, L-733,060 (1x, 0.0176 μg/μl, n = 8; 10x, 0.176 μg/μl, n = 7), or vehicle during period of large irregular hippocampal field activity (LIA). (C) Microinjection of L-733,060 into MS attenuated formalin-induced suppression of CA1 PS. The PS traces on the left and histograms are built as explained in Fig. 1. (D) Histograms illustrating lack of effect of L-733,060 pre-treatment on fast Fourier transform (FFT; resolution of 0.5 Hz) parameters of hippocampal theta wave activity (FFT theta peak power, left and FFT theta peak frequency, right). FFT analysis was performed on theta wave activity recorded in the first 5 min after formalin injection. The FFT theta peak power was normalised to spontaneous theta power recorded prior to the experiment. Data presented are mean ± S.E.M. Significant difference (p < 0.05): * vs. Vehicle MS; two-tailed unpaired t-test.

Since the time course of effect of 1x L-733,060 (n = 8) and 10x L-733,060 (n = 7) on formalin response overlapped (Treatment, F_1, 299_ = 0.29, p > 0.5; two-way RM ANOVA; data not shown), the two groups were combined and compared with the control response. In this context, twoway RM ANOVA showed a significant effect of pre-treatment (Treatment, F_1, 483_ = 26.26, p < 0.0001; data not shown), while Bonferroni post-hoc test showed that suppression of PS amplitude was attenuated with L-733,060 pre-treatment at all time-points except at 1^st^ and 7^th^ min following formalin injection. Consistently, the average percentage of PS amplitude was significantly higher in L-733,060 pre-treated animals as compared to vehicle pre-treated control group (Figure 3C; p < 0.0001; two-tailed unpaired t-test) suggesting that the suppression of CA1 PS evoked on formalin injection was attenuated on pre-treatment with L-733,060. The control amplitudes of the PS in the two minutes preceding injection of formalin were not different in the vehicle pre-treatment group vs. the L-733,060 pre-treatment group (6.16 ± 0.32 mV vs. 6.23 ± 0.25 mV, p > 0.8; two-tailed unpaired t-test; data not shown).

### Theta wave activity

#### Effect of microinjection of L-733,060 on hippocampal theta wave activity

Hippocampal theta field activity was recorded concurrently with PS in the preceding experiments. Theta reflects synchronized oscillations of hippocampal neurons during processing of information and is mediated, in part by the medial septum. However, microinjection of SP into MS evoked a weak theta activation that was not clearly separated from other treatmetn groups in the pilot study. In this context, the average duration of theta (s/min) in five minutes following various pre-treatments in the pilot study, namely Vehicle vs. SP (2 μg/μl) in LS vs. SP (1 μg/μl) in MS vs. SP (2 μg/μl) in MS, were 1.49 ± 0.76 (n=5) vs. 2.91 ± 0.85 (n=7) vs. 4.64 ± 1.47 (n=7) vs. 8.26 ± 2.16 (n=9), respectively. Analysis revealed a lack of significant difference among the groups (Groups, p > 0.1; Kruskal-Wallis test). In the larger sample that included the response to SP1 (2 μg/μl) in the vehicle and the L-733,060 pre-treatment groups, the average duration of theta (s/min) within first 5 min was also low (7.44 ± 2.16, n = 20) that is comparable to that seen in the pilot study.

On the other hand, a relatively robust increase in average duration of theta (s/min) was seen in the five minutes afer C1 microinjection into medial septum (26.74 ± 2.95, n = 10) or on hind paw injecion of formalin (26.88 ± 2.29, n = 9). Therefore, the effect of L-733,060 on FFT theta peak frequency and FFT theta peak power was analyzed on the background of robust theta seen in the 5 min after carbachol or formalin injection. To note that the data presented in the foregoing section showed that L-733,060 attenuated formalin-induced suppression suggesting that transmission at NK1R played a role in formalin induced septo-hippocampal responses.

Statistical analysis revealed a lack of significant effect of L-733,060 pre-treatment on carbachol-induced FFT theta peak frequency (Groups, F_2, 12_ = 1.70, p > 0.2, n = 5; one-way ANOVA), FFT theta peak power (Groups, F_2, 12_ = 0.57, p > 0.5, n = 5; one-way ANOVA) and average duration of theta per minute (Groups, F_2, 12_ = 0.04, p > 0.9, n = 5; one-way ANOVA). In this context, the FFT theta peak frequencies (Hz) with C1, L-733,060 + C2 and C3 were 4.94 ± 0.16, 4.50 ± 0.26 and 4.32 ± 0.30, respectively. The corresponding values of normalized FFT theta power were 0.42 ± 0.06, 0.53 ± 0.07, and 0.51 ± 0.09. To note that the drug-induced FFT theta peak power was normalized to spontaneous theta. The values of theta duration (s/min) corresponding to the three repeat microinjections were 21.73 ± 3.93 vs. 21.42 ± 7.20 and 18.87 ± 6.94.

Similarly, vehicle pre-treatment also did not affect FFT theta peak frequency (Groups, F_2, 12_ = 1.12, p > 0.3, n = 5; one-way ANOVA), FFT theta peak power (Groups, F_2, 12_ = 0.92, p > 0.4, n = 5; one-way ANOVA) and average duration of theta (s/min) (Groups, F_2, 12_ = 1.33, p > 0.3, n = 5; one-way ANOVA). The FFT theta peak frequencies (Hz) corresponding to C1, vehicle + C2 and C3 were 5.16 ± 0.26, 4.82 ± 0.25 and 4.72 ± 0.13. The normalized FFT theta peak power with the three repeat microinjections were 0.48 ± 0.06, 0.62 ± 0.11, and 0.66 ± 0.12 while the corresponding durations were 31.74 ± 3.34 vs. 33.83 ± 6.57 and 32.79 ± 7.32.

Since the FFT theta peak power was normalized to spontaneous theta, we also compared the theta parameters of spontaneous theta recorded at the beginning of C1. The FFT theta peak power of the spontaneous theta in the group pre-treated with vehicle (n = 5) or L-733,060 (n = 5) was not different from each other (0.05 ± 0.01 mV^2^ vs. 0.04 ± 0.01 mV^2^, p > 0.5; two-tailed unpaired t-test). The FFT theta peak frequency of the spontaneous theta in the group pretreated with vehicle (n = 5) or L-733,060 (n = 5) was also not different from each other (4.20 ± 0.20 Hz vs. 3.80 ± 0.10 Hz, p > 0.5; two-tailed unpaired t-test).

With regards to formalin test, pre-treatment with L-733,060 (1x and 10x) significantly attenuated average duration of theta (s/min) in the first 5 min following injection of formalin (‘Vehicle MS’ group vs. ‘L-733,060 MS’ group being 26.88 ± 2.29 (n = 9) vs. 11.82 ± 1.91 (n = 15); p < 0.0001; two-tailed unpaired t-test). However, normalised FFT theta peak power (Figure 3D left; p > 0.4; two-tailed unpaired t-test) and FFT theta peak frequency (Figure 3D right; p > 0.6; two-tailed unpaired t-test) were unaffected. The FFT theta peak power of the spontaneous theta in the group pre-treated with vehicle (n = 9) or L-733,060 (n = 15) was not different from each other (0.07 ± 0.01 mV^2^ vs. 0.05 ± 0.006 mV^2^, p > 0.05; two-tailed unpaired t-test).

### Effect of intraseptal drugs on animal behaviours and theta activation

#### General

The observer was blinded to the pharmacological treatments in the experiments described below. Acute behaviours monitored included ambulation in the novel open field and the formalin test, and the formalin-induced nociceptive licking and flinching.

L-733,060 was administered as pre-treatment, which is consistent with the protocol in anaesthetized animal. The antagonist sufficed to antagonize both intraseptal SP- and formalin injection-induced hippocampal responses in anaesthetized animal (see above). On the other hand, SST was microinjected just prior to behavioural experiment to minimize the loss of effect due to diffusion of the agent over time. This precaution was taken since the pattern of effect of SST was not characterized as was done for L-733,060. The concentration of SST (3.38 μg/μl or 6.76 μg/μl) was selected based on published work where it has been shown to disrupt theta in exploratory behaviour (Bassant *et al*., 2005).

### Behaviour

#### Effect of microinjection of the NK1R receptor antagonist, L-733,060, on exploration in novel open field

L-733,060 (1x, 0.0176 μg/μl; 0.5 μl) or vehicle was microinjected into the MS or the LS (Figure 4A) 15 min before the open field test. Normally, the animals’ activity in the novel open field chamber is highest within the first 15 minutes of exposure to the test chamber (Ang *et al*., 2015). Thus, the effect of pre-treatment with L-733,060 was analysed in this time frame. This time of observation is characterized by an initial high ambulation (rising phase, 1-5 min), followed by declining ambulation (declining phase, 6-10 min) and finally habituation where the animal, more or less, comes to rest (resting phase, 11-15 min). Statistical analysis showed a lack of significant effect of L-733,060 treatment on ambulation (Figure 4B, left; Treatment, F_2, 60_ = 1.74, p > 0.1, two-way RM ANOVA). However, a significant effect of time was noted (Figure 4B, left; Time, F_2, 60_ = 78.03, p < 0.0001, two-way RM ANOVA). To get a better measure of effect of time, the group means at individual phases were further analysed. The ambulation in the three groups was not different in the rising phases (Groups, F_2, 20_ = 0.21, p > 0.8, one-way ANOVA; data not shown). However, microinjection of L-733,060 into MS significantly decreased ambulation in the declining and resting phases (Groups, p < 0.02 at least; Kruskal-Wallis test followed by Dunn’s post-hoc test; data not shown) suggesting the basis of the time-specific effects among groups. The pattern of decrease was similar with microinjection into MS and LS, though only the former reached significance vis-à-vis vehicle pre-treated animals.

**Figure 4.**
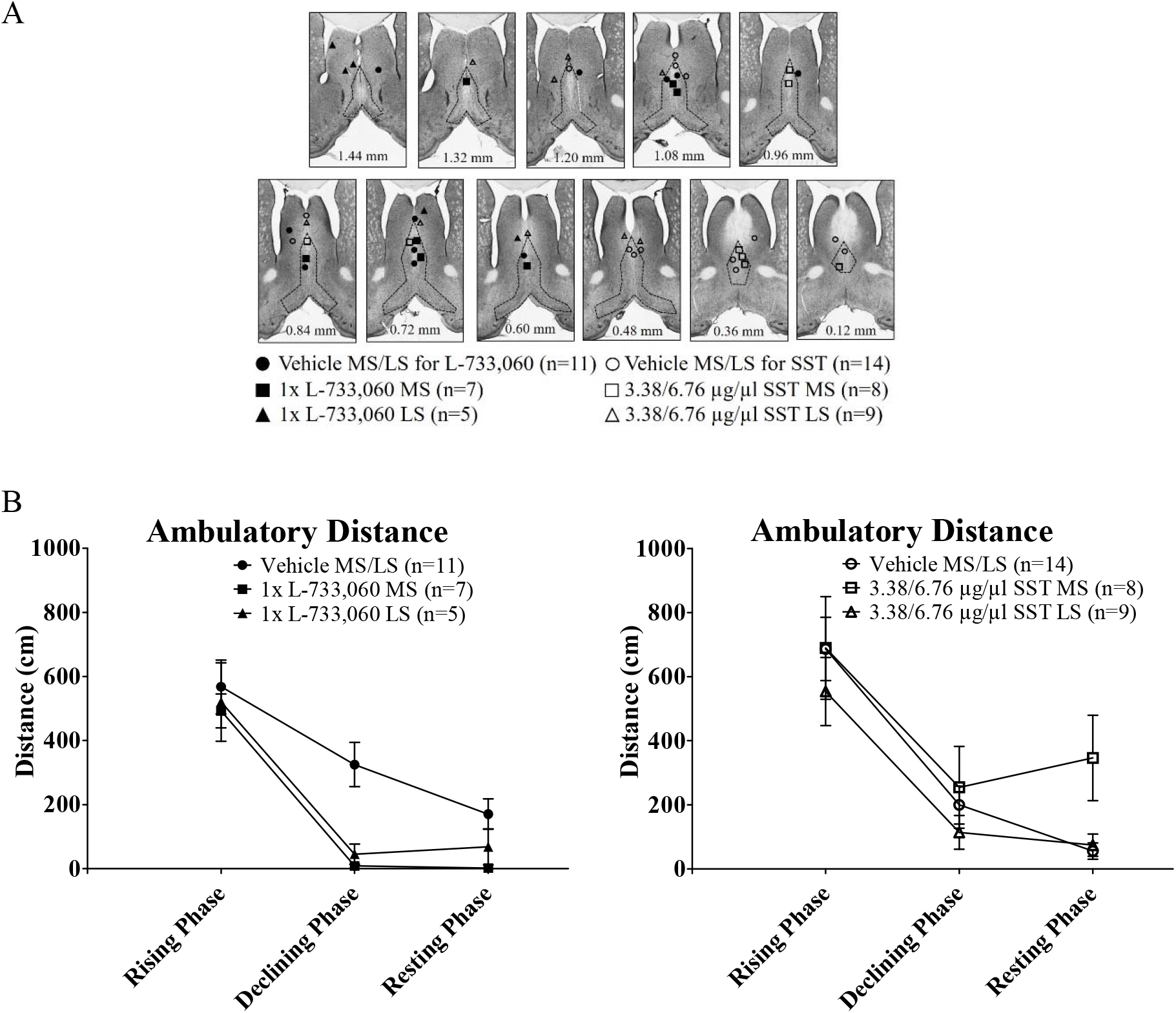
Intraseptal L-733,060 and somatostatin (SST) differentially modulates exploratory behaviour in a novel open field test. (A) Composite representation of microinjection sites in the medial septum (MS) and lateral septum (LS). The animal was placed into the novel open field chamber 15 min following intraseptal microinjection of L-733,060 (1x, 0.0176 μg/μl, MS n=7, LS n=5) or the corresponding vehicle, or immediately after microinjection of SST (3.38 μg/μl, MS n=3, LS n=5; 6.76 μg/μl, MS n=5, LS n=4) or the corresponding vehicle. The rising phase, declining phase and resting phase correspond to 1-5 min, 6-10 min and 11-15 min following exposure to the novel open field chamber. (B) Time course graphs of ambulatory distance in block of 5 mins in the first 15 minutes upon exposure to the novel open field chamber. Although the effect of treatment is insignificant, there is a trend towards decreasing ambulation in the declining phase upon microinjection of L-733,060 (left) into the MS and LS, and increasing ambulation in the resting phase with microinjection of SST (right) into the MS. Data are expressed as mean + S.E.M.

The decrease in ambulation was very robust such that ambulatory movement of only one or two animals pre-treated with L-733,060 into MS or LS reached threshold for calculating speed (i.e. animal movement for more than 2s) in the 6-15^th^ min period. As a result, speed was analysed only during the rising phase. In the rising phase, the speed (cm/s) for the Vehicle MS/LS group vs. 1x L-733,060 MS group vs. 1x L-733,060 LS group were: 103.7 ± 27.31 (n = 11) vs. 59.11 ± 14.34 (n = 7) vs. 59.46 ±11.31 (n = 5). The speed was not different (Groups, F_2, 20_ = 1.22, p > 0.3, one-way ANOVA; data not shown), although the speed following L-733,060 pre-treatment tended to be on the lower side.

#### Effect of microinjection of SST on exploration in novel open field

In separate set of experiments, SST (3.38 μg/μl, n = 3 in MS and 5 in LS; 6.76 μg/μl, n = 5 in MS and 4 in LS; 0.5 μl) was microinjected right before exposure to the novel open field chamber. Because of overlapping effects (data not shown), the two doses for a given site were combined. The three groups thus analysed for effect of SST were Vehicle MS/LS group, 3.38/6.76 μg/μl SST MS group and 3.38/6.76 μg/μl SST LS group (Figure 4). Statistical analysis revealed a lack of significant effect of SST treatment (Figure 4B, right; Treatment, F_2, 56_ = 1.99, p > 0.1; two-way RM ANOVA). However, a significant effect of time was observed (Figure 4B, right; Time, F_2, 56_ = 30.64, p < 0.0001). To get a better measure of effect of time, the means of various groups at selected time points were further analysed for differences. The time points selected for analysis corresponded to the rising phase, the declining phase and the resting phase of the open field (Figure 4). The ambulation in the three groups was not different in the rising phase and declining phases (Groups, F_2, 29_ = 0.62, p > 0.5 at least; one-way ANOVA). However, microinjection of SST into MS, but not LS significantly increased ambulation in resting phase (11-15min; Groups, p < 0.05; Kruskal-Wallis test followed by Dunn’s post-hoc test) pointing to the basis of the time-specific effect among groups.

In order to get a better measure of the effect of SST on ambulation in the resting phase of the open field, animals that showed no measurable ambulation (i.e. zero ambulation) were excluded and the rest compared for an effect of SST on ambulation in the resting phase. The number of animals that were so excluded were 6 of 14 (42.9%), 1 of 8 (11.1%) and 2 of 9 (22.2%) for the Vehicle MS/LS group, 3.38/6.76 μg/μl SST MS group and 3.38/6.76 μg/μl SST LS group, respectively. The distance (cm) moved by the remaining animals in the three groups, i.e. the Vehicle MS/LS group, 3.38/6.76 μg/μl SST MS group and 3.38/6.76 μg/μl SST LS group was 98.14 ± 39.21 (n = 8), 395.80 ± 142.80 (n = 7) and 95.75 ± 41.89 (n = 7), respectively. Since the ambulation in Vehicle MS/LS group and 3.38/6.76 μg/μl SST LS group was similar (p > 0.9 two-tailed unpaired t-test), the two were combined into a composite Control group (ambulation 97.02 ± 27.59 cm, n = 15). The ambulation in the composite Control group was significantly lower than in 3.38/6.76 μg/μl SST MS group (p < 0.009, two-tailed unpaired t-test).

As a comparison, we also analyzed the changes during period of high ambulation (i.e. rising phase in open field) in the foregoing fashion. In this period, none of the animals in the composite Control group and the 3.38/6.76 μg/μl SST MS group exhibited zero ambulation. The distance (cm) moved by the animals in the two groups, i.e. composite Control group and 3.38/6.76 μg/μl SST MS group was 634.80 ± 72.96 (n = 23) and 689.90 ± 160.30 (n = 8), respectively. The ambulation was similar among the two groups (p > 0.7, two-tailed unpaired t-test).

The speed of ambulation was analysed at the two ends of the behaviour, namely the peak of animal ambulation during the rising phase and at the resting phase. However, to note, that ambulation in number of animals did not reach threshold for measurement of speed, especially after the rising phase. In the rising phase, the speed (cm/s) for Vehicle MS/LS group vs. 3.38/6.76 μg/μl SST MS group vs. 3.38/6.76 μg/μl SST LS group were: 40.57 ± 2.39 (n = 14) vs. 40.29 ± 3.67 (n =7) vs. 57.04 ±11.12 (n =9). The speed was not different (Groups, p > 0.8, Kruskal-Wallis test; data not shown).

Likewise, for the resting phase the values of speed (cm/s) were, in order: 50.72 ± 20.09 (n = 4) vs. 32.03 ± 2.60 (n = 6) vs. 32.10 ± 3.38 (n = 4). These were not significantly different (Groups, p > 0.7, Kruskal-Wallis test; data not shown).

#### Effect of microinjection of the NK1R receptor antagonist, L-733,060, on formalin-induced nociceptive behaviours

Formalin (0.1 ml, 1.25%) injection into the hind paw evoked a biphasic increase in animal agitation marked by increase in ambulation, nociceptive licking and flinching similar to that reported before (Ang *et al*., 2015; Lee *et al*., 2011; Tai *et al*., 2006). The biphasic pattern is characterized by a peak of behavioural responses in the first 5 min after formalin injection or Phase 1 (1-5 min) and another peak in Phase 2 (Phase 2 peak; 16-20 min). The Phase 2 *per se* corresponds to 11^th^ to 60^th^ min following injection of formalin. The interphase (6^th^ to 10^th^ min after formalin injection) is marked by quiescence with relatively low nociceptive and ambulatory activity (Ang *et al*., 2015; Lee *et al*., 2011; Tai *et al*., 2006).

The effect of L-733,060 on formalin-induced nociceptive behaviours was examined at two doses of the antagonist (Figure 5A: 1x, 0.0176 μg/μl, n= 9 in MS and 14 in LS, and 10x, 0.176 μg/μl, n=7 in MS; 0.5 μl). The time course of change for each of the formalin-induced nociceptive behaviours-i.e. licking and flinching-was compared between the two pretreatment groups. Two-way RM ANOVA showed that a given formalin-induced behaviour following microinjection of low or high dose of L-733,060 into MS was similar between the two groups (Treatment, F_1, 154_ = 0.98 p > 0.3 at least). Thus, the two pre-treatment groups were consolidated as a ‘L-733,060 MS’ group (n = 16). The ‘L-733,060 LS’ group was microinjected with only the low dose of L-733-060 into LS (n = 14). The formalin-induced responses among animals microinjected with vehicle into MS (n = 6) or the LS (n = 4) were also similar (Treatment, F_1, 88_ = 0.06 p > 0.8 at least, two-way RM ANOVA). Thus, two groups were also combined (‘Vehicle MS/LS’ group, n = 10).

**Figure 5.**
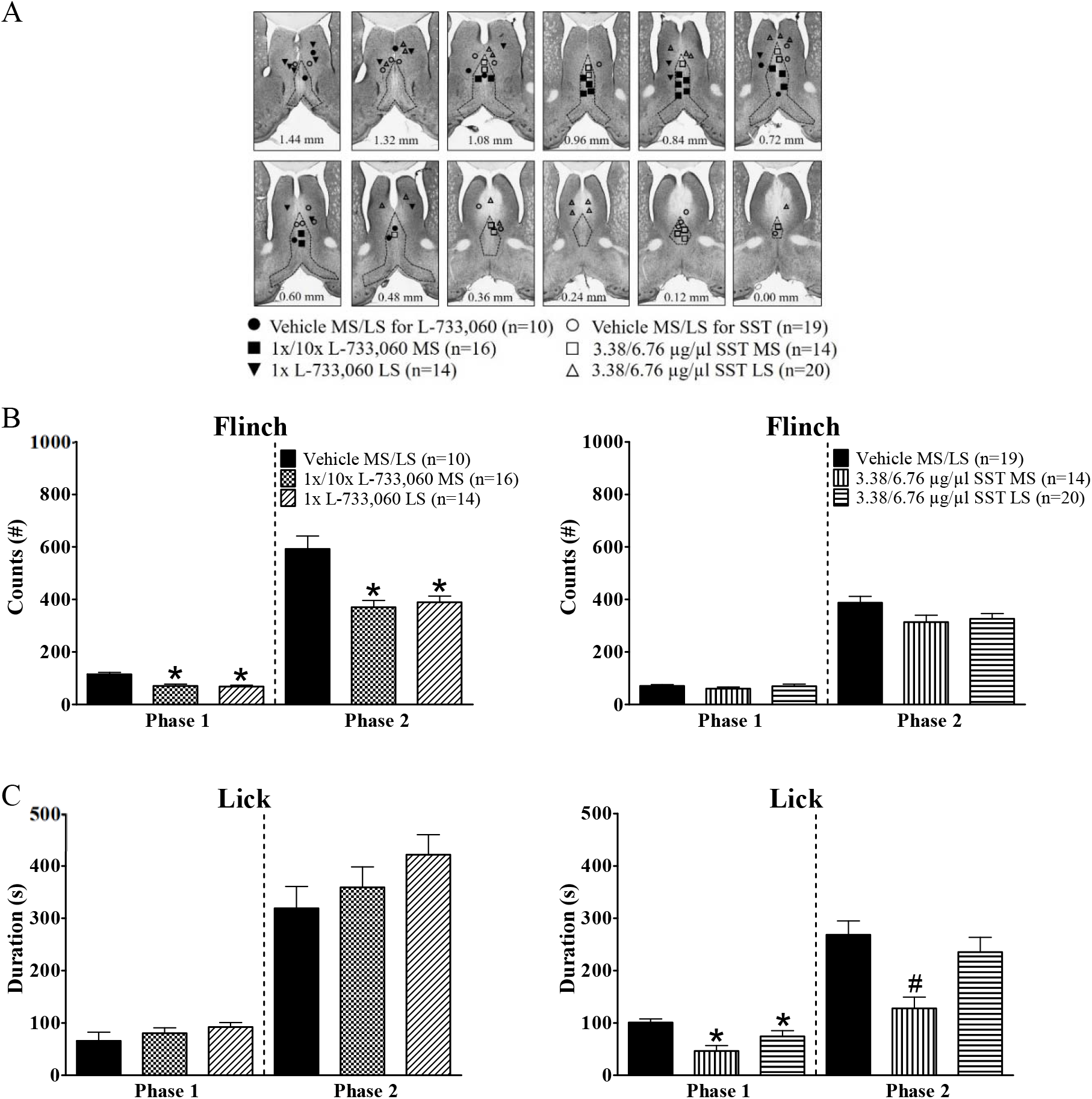
Intraseptal L-733,060 and somatostatin (SST) differentially modulate nociceptive behaviours in the formalin test. (A) Composite representation of microinjection sites in the medial septum (MS) and lateral septum (LS). Formalin (1.25%, 0.1ml) was injected into right hind paw either 15 min (left) or immediately after microinjection (right) of L-733,060 (1x, 0.0176 μg/μl, MS n=9, LS n=14; 10x, 0.176 μg/μl, MS n = 7) or SST (3.38 μg/μl, MS n=7, LS n=10; 6.76 μg/μl, MS n=7, LS n=10) or vehicle. (B) Histograms illustrating the cumulative number of flinches of the right hind paw in Phase 1 (first 5 min) and Phase 2 (11-60 min) of the formalin test on intraseptal microinjection of either L-733,060 (left) or SST (right). Intraseptal microinjection of L-733,060 but not SST attenuated flinching behaviour in both phases. (C) Histograms illustrating the cumulative duration of licking of the right hind paw in Phase 1 (first 5 min) and Phase 2 (11-60 min) of the formalin test on intraseptal microinjection of either L-733,060 (left) or SST (right). The parameter was recorded in the same experiments as in B. Intraseptal SST, but not L-733,060 significantly attenuated licking behaviours in both phases. Data are mean ± S.E.M. Significant difference (p < 0.05): (B, C): * vs. ‘Vehicle MS/LS’ group, # vs. ‘Vehicle MS/LS’ and ‘SST LS’ groups; one-way ANOVA followed by Newman-Keuls post-hoc test.

Compared to Vehicle, microinjection of L-733,060 into both MS and LS attenuated formalin-induced flinching at various time points along its time course (Treatment, F_2, 407_ = 17.92, p < 0.0001; two-way RM ANOVA followed by Bonferroni post-hoc test; data not shown). Whereas, licking was not significantly affected (Treatment, F_2, 407_ = 2.28, p > 0.1; two-way RM ANOVA; data not shown).

Consistently, phase analysis indicated that pre-treatment with L-733,060 signifcantly reduced flinching in both Phase 1 and Phase 2 of the formalin test (Figure 5B, left; Phase 1, Groups, F_2, 37_ = 13.21, p < 0.0001; Phase 2, Groups, F_2, 37_ = 13.46, p < 0.0001; one-way ANOVA followed by Newman-Keuls post-hoc test). Whereas, licking was not affected across both phases of the formalin test (Figure 5C, left; Phase 1, Groups, F_2, 37_ = 1.16, p > 0.3; Phase 2, Groups, F_2, 37_ = 1.52, p > 0.2; one-way ANOVA).

#### Effect of microinjection of SST on formalin-induced behaviours

In separate set of experiments, SST (3.38 μg/μl, n = 7 in MS and 10 in LS; 6.76 μg/μl, n = 7 in MS and 10 in LS; 0.5 μl) was microinjected just prior to right hind paw formalin injection (0.1 ml, 1.25%). The data is reported separately from above since these experiments were monitored independently from the foregoing by another observer.

Two-way RM ANOVA showed that a given formalin-induced behaviour following microinjection of low or high dose of SST into MS was similar between the two groups (Treatment, F_1, 132_ = 0.32, p > 0.5 at least). Thus, the two groups were combined into a ‘3.38/6.76 μg/μl SST MS’ group (n=14). Likewise, statistics also revealed a similar patterns of responses on microinjection of the two doses into LS (Treatment, F_1, 306_ = 2.73, p > 0.1 at least). The two doses were combined, the combined group being termed ‘3.38/6.76 μg/μl SST LS’ group (n=20). Control data obtained on microinjection of vehicle into the MS or the LS were also collapsed since the patterns of responses were similar among the two groups (‘Vehicle MS/LS’ group, n=19; Treatment, F_1, 198_ = 0.97, p > 0.3 at least).

Unlike L-733,060, microinjection of SST into MS significantly reduced licking up to the 25^th^ min of the formalin-induced time course (Treatment, F_2, 550_ = 8.10, p < 0.001; two-way RM ANOVA with Bonferroni post-hoc test; data not shown). Some decrease in licking was observed on microinjection into LS but individual time points did not reach significance with the statistical test (data not shown).

Whereas, nociceptive flinching was not different in SST treated animals, as compared to control, across the individual time points of the time course, although an overall effect of treatment was observed on this behaviour (Treatment, F_2, 550_ = 3.44, p < 0.04; two-way RM ANOVA followed by Bonferroni post-hoc test; data not shown).

Consistently, phase analysis showed that intraseptal microinjection of SST reduced licking (Figure 5C, right; Phase 1, Groups, F_2, 50_ = 7.32, p < 0.002; Phase 2, Groups, F_2, 50_ = 6.77, p < 0.003; one-way ANOVA followed by Newman-Keuls post-hoc test) but without affecting flinching (Figure 5B, right; Phase 1, Groups, p > 0.2, Kruskal-Wallis test; Phase 2, Groups, F_2, 50_ = 2.94, p > 0.06, and one-way ANOVA). The effect of intraseptal SST on licking was most profound in 3.38/6.76 μg/μl SST MS group, attenuating licking in both Phase 1 and Phase 2, while licking was attenuated only in Phase 1 in the 3.38/6.76 μg/μl SST LS group.

#### Effect of intraseptal microinjection on formalin-induced ambulation

Microinjection of L-733,060 into MS or LS did not affect ambulation (Figure 6, left; Treatment, F_2, 111_ = 3.00, p > 0.06). Ambulation at a given phase of behaviour did not always reach threshold for measurement of speed in a number of animals. Thus, in context of ambulatory speed, the group sizes are uneven and the phases were analysed separately. In relation to Phase 1, the ambulatory speed (cm/s) for Vehicle MS/LS vs. 1x/10x MS L-733,060 vs. 1x LS L-733,060 were 178.90 ± 17.20 (n = 10) vs. 189 ± 29 (n = 16) vs. 215 ± 46.30 (n = 14). The speeds were not statistically different (Groups, F_2, 37_ = 0.06 p > 0.9; one-way ANOVA). The speeds were also not statistically different at interphase (p > 0.9; Kruskal-Wallis test) and the peak of Phase 2 (Groups, F_2, 33_ = 1 p > 0.3; one-way ANOVA). At interphase, the speeds (cm/s) for the three groups were, in order, 136.40 ± 42.40 (n = 4) vs. 149.40 ± 22.50 (n = 3) vs. 147.40 ± 30.60 (n = 8), while that at peak of Phase 2 were 140.30 ± 12.40 (n = 9) vs. 121.20 ± 6.50 (n = 14) vs. 135.60 ± 11.60 (n = 13).

**Figure 6.**
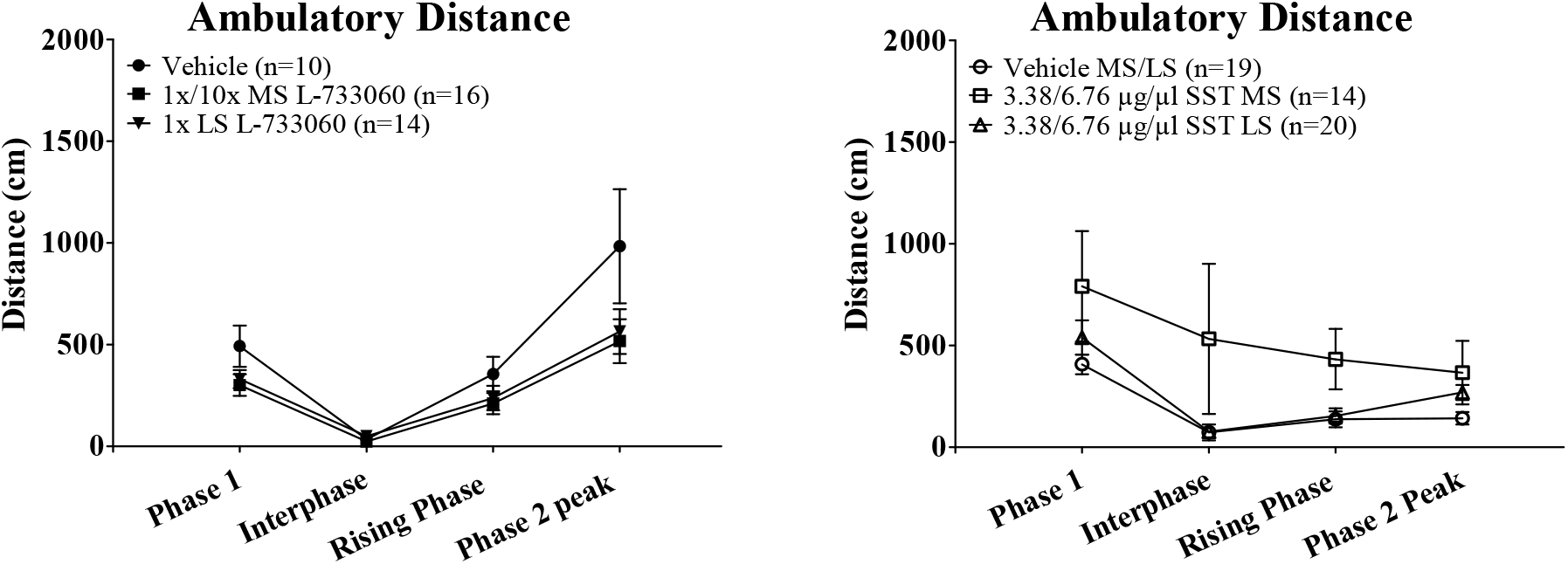
Intraseptal microinjection of somatostatin (SST), but not L-733,060 affects formalin-induced agitation. Formalin (1.25%, 0.1ml) was injected into the right hind paw 15 minutes following L-733,060 microinjection and immediately after SST. The time course graphs are built as explained in Fig. 4. While, L-733,060 did not significantly affect ambulation, SST pre-treatment evoked an overall significant effect on ambulatory distance.

On the toher hand, two-way ANOVA followed by Bonferroni post-hoc test revealed an overall significant effect of SST treatment on ambulation (Figure 6, right; Treatment, F_2, 150_ = 3.44, p < 0.04). However, the individual time points were not significantly different, though a trend towards increase in ambulation was observed at interphase. As above, ambulatory speed at different phases was analysed separately. In relation to Phase 1, the ambulatory speeds (cm/s) for Vehicle MS/LS vs. 3.38/6.76 μg/μl SST MS vs. 3.38/6.76 μg/μl SST LS were 122.30 ± 10.10 (n= 19) vs. 149.80 ± 16 (n=14) vs. 134.40 ± 7.50 (n=20). The speeds were not statistically different (Groups, F_2, 50_ = 1.49, p > 0.2; one-way ANOVA). The speeds were also not statistically different at interphase (p > 0.1; Kruskal-Wallis test) and Phase 2 peak (Groups, F_2, 42_ = 0.03, p > 0.9; one-way ANOVA). At interphase, the speeds (cm/s) for the three groups were, in order, 107.80 ± 15.20 (n = 3) vs. 121.30 ± 11.50 (n = 7) vs. 141.90 ± 16.40 (n = 8), while that at peak of Phase 2 were 140.50 ± 19.30 (n = 14) vs. 135 ± 18.90 (n=12) vs. 137.70 ± 9.10 (n = 19).

Interestingly, animal ambulation at interphase showed a wide range, varying from zero to higher values. In this context, the number of animals showing zero ambulation across different groups was as follows: Vehicle MS/LS vs. 3.38/6.76 μg/μl SST MS vs. 3.38/6.76 μg/μl SST LS-11 of 19 vs. 5 of 14 vs. 9 of 20. As with open field, animals showing zero ambulation were excluded and the rest compared for an effect of SST on ambulation in the interphase. The distance (cm) covered by ambulatory animals during interphase in Vehicle MS/LS vs. 3.38/6.76 μg/μl SST MS vs. 3.38/6.76 μg/μl SST LS was 170.40 ± 85.20 (n = 8) vs. 827.70 ± 560.30 (n = 9) vs. 136.80 ± 38.80 (n = 11). The ambulation of Vehicle MS/LS vehicle and 3.38/6.76 μg/μl SST LS groups was similar (p > 0.6 two-tailed unpaired t-test). These two groups were combined together to form a composite Control group (ambulation: 151 ± 41.10 cm, n = 19). A comparison between the animals in the composite Control vs. 3.38/6.76 μg/μl SST MS showed that the ambulation in the latter was significantly higher (p < 0.04, two-tailed unpaired t-test).

In contrast, during period of high ambulation in phase 1, where none showed zero ambulation, the ambulations in composite Control group of animals (474.50 ± 49.80 cm, n = 39) and the 3.38/6.76 μg/μl SST MS group (790.70 ± 271.20 cm, n = 14) were not different from each other (p > 0.7, two-tailed unpaired t-test).

### Theta wave activity

#### General

Theta wave activity from hippocampal stratum radiatum and stratum lacunosum-moleculare, concomitantly monitored with behaviours in the open field and the formalin test, were analysed using FFT (0.5Hz resolution). However, the number of experiments with successful recording was less than for behaviour. As a result, the sample size for analysis of field activity is somewhat lower than for behaviour. At an extreme, very few animals’, especially control animals, showed clear theta wave activity in the study investigating the effect of SST on animal behaviour in novel open field. Thus, theta wave activity was not analysed here.

Further, the behavioural effect of L-733,060 was clear in the first 10 min of open field exploration that covered the rising and declining phase of exploration. Thus, we analysed the two time-blocks in this period for L-733,060-induced change in theta parameters. However, the number of animals displaying clear theta declined from the rising to the declining phase of open field. As a result, the sample size is uneven from one time-block to the other and each phase was analysed separately.

The effects of the drug in the formalin test, however, were on a more consistent background of theta activation. In the formalin test, the theta was analysed for the following phases: Phase 1 (1-5 min), interphase (6-10 min), rising phase (11-15 min) and Phase 2 peak (16-20 min). The Phase 1 and Phase 2 peak correspond to peak of behavioural activity, while the interphase is a period of relative behavioural quiescence. The rising phase of the formalin test reflects an increasing nociception from the low of interphase towards the peak at Phase 2.

#### Comparison of theta in novel open field vs. formalin test

The relative power of theta in open field vis-à-vis the formalin test was analysed in control, vehicle treated animals. In this context, the normalised powers in first 10min of novel open field vs. Phase 1 vs. Phase 2 of the formalin test were 1.12 ± 0.07 (n = 7) vs. 0.50 ± 0.05 (n = 15) vs. 0.40 ± 0.02 (n = 15). The powers during Phase 1 and Phase 2 was significantly lower as compared to that in the open field (Groups, F_2, 34_ = 37.17, p < 0.0001; one-way ANOVA followed by Newman-Keuls test). Thus, the drug effects in the formalin test were against a lower power of theta wave activity. Since the power was normalised to the power of theta wave activity recorded during the exploration of a known environment (i.e. control theta), we compared the power of control theta between the two set of experiments. The power of control theta was similar in animals used in test of exploration in novel open field vs. animals used for the formalin test (0.1144 ± 0.0361 mV^2^ (n = 7) vs. 0.0963 ± 0.0122 mV^2^ (n = 15), p > 0.5; two-tailed unpaired t-test).

Unlike power, the difference in FFT theta peak frequency of theta wave recorded in vehicle pre-treated animals during open field and the formalin test were not that clearly separated. Thus, the FFT theta peak frequency in the first 10min of the open field and Phase 1 of the formalin test were similar though significantly higher than that in Phase 2 of the formalin test (Groups, F_2, 34_ = 15.75, p < 0.0001; one-way ANOVA followed by Newman-Keuls test). The FFT theta peak frequency (Hz) in novel open field vs. Phase 1 vs. Phase 2 of the formalin test were 7.32 ± 0.07 (n = 7) vs. 7.37 ± 0.09 (15) vs. 6.74 ± 0.09 (n = 15).

#### Effect of microinjection of L-733,060 on theta activation in novel open field

Intraseptal L-733,060 did not significantly affect theta power and frequency in the open field. In this context, the power during the rising phase of open field in Vehicle MS/LS group vs. 1x L-733,060 MS vs. 1x L-733,060 LS was 1.21 ± 0.09 (n = 7) vs. 0.93 ± 0.21 (n = 5) vs. 1.39 ± 0.13 (n = 5). The values were not different from each other (Groups, F_2, 14_ = 2.36, p > 0.1, one-way ANOVA). The corresponding values in the declining phase were 1.04 ± 0.07 (n = 6) vs. 0.57 + 0.19 (n = 4) vs. 1 ± 0.15 (n = 4). These are also not different from each other (Groups, p > 0.1, Kruskal-Wallis test).

The frequency of the theta wave activity was similar across the treatment groups (rising phase: Groups, p > 0.3, Kruskal-Wallis test; declining phase: Groups, p > 0.3; Kruskal-Wallis test; data not shown).

#### Effect of microinjection of L-733,060 on formalin-induced theta activation

We also examined the effect of microinjection of L-733,060 on duration of formalin-induced theta since this parameter was affected by the treatment in anaesthetized animal. The microinjection of L-733,060 into MS or LS of behaving animal did not affect the duration of formalin-induced theta from 1-20min (i.e. from Phase 1 to the peak of Phase 2; Treatment, F_2, 75_ = 1.12, p > 0.3; twoway RM ANOVA; data not shown).

However, microinjection of L-733,060 into MS, but not LS, evoked a parallel shift in power with a signficant reduction of power in Phase 1, interphase and rising phase after formalin test (Figure 7A, left; Treatment, F_2, 75_ = 9.07, p < 0.002; two-way RM ANOVA followed by Bonferroni post-hoc test). Furthermore, the average of normalized theta power in the four phases was significantly lower (Groups, F_2, 25_ = 9.07, p < 0.002; one-way ANOVA followed by Newman-Keuls post-hoc test) in the 1x/10x L-733,060 MS group (0.28 ± 0.03, n = 11) vs. Vehicle MS/LS group (0.47 ± 0.04, n = 6) and 1x L-733,060 LS group (0.41 ± 0.03, n= 11).

**Figure 7.**
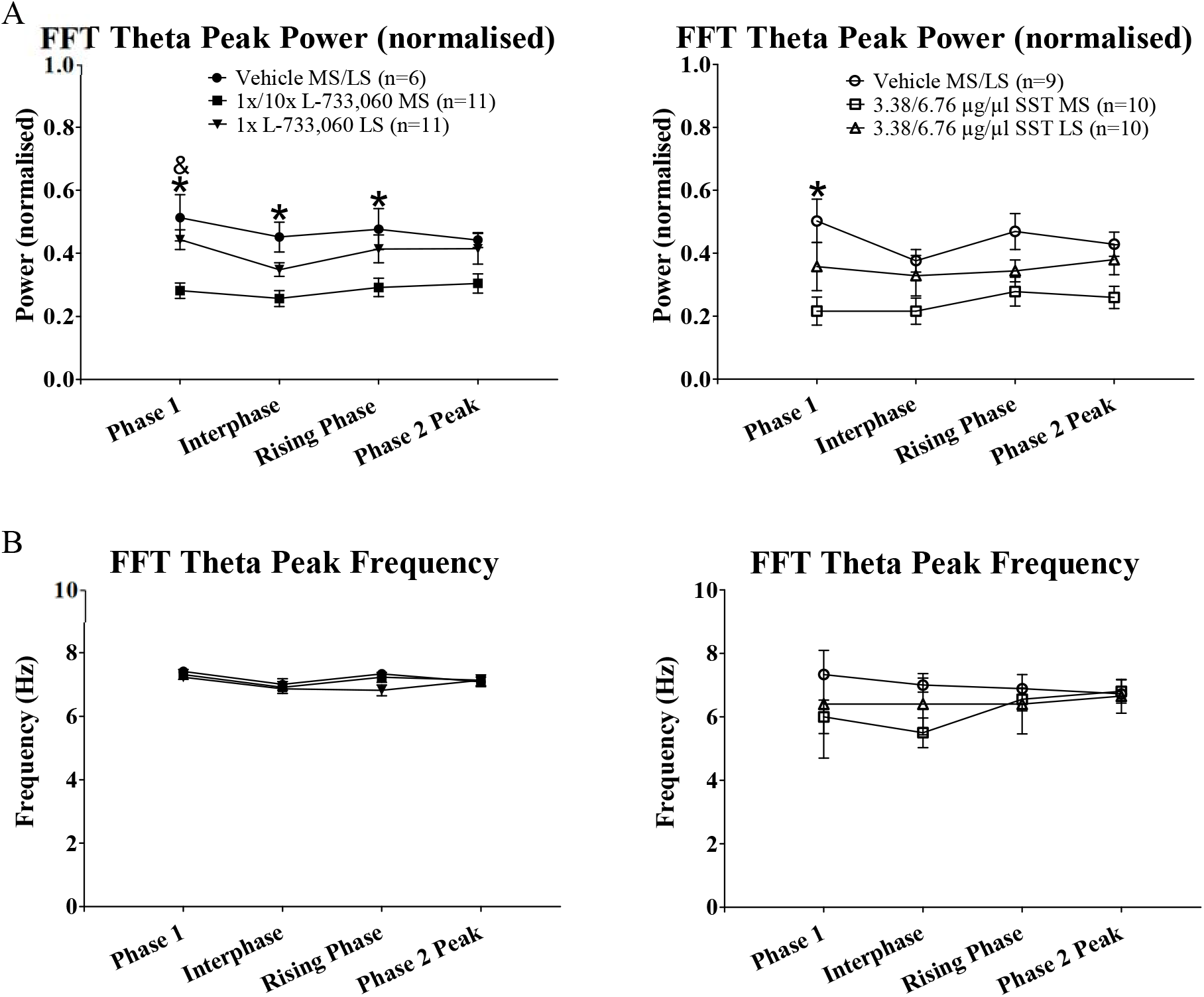
Intraseptal microinjection of L-733,060 and somatostatin (SST) attenuated formalin-induced theta activity. The hippocampal theta wave activity was induced on hind paw injection of formalin (1.25%, 0.1ml). L-733,060 was microinjected 15 min before formalin injection while SST was microinjected immediately before. The time of injection of formalin was taken as 0 min. Hippocampal theta segments of at least 2s duration were analysed using Fast Fourier Transform (FFT; resolution of 0.5Hz). The FFT theta peak power and the corresponding frequency were averaged in blocks of 5 min. Both parameters are presented at 4 time points of the formalin test (Phase 1: first 5 min, Interphase: 6-10 min, Rising Phase: 1115 min, Phase 2 Peak: 16-20 min). (A) Time course of FFT theta peak power in blocks of 5 min. FFT theta peak power was normalised to exploratory FFT theta peak power recorded prior to the experiment. Note that both drugs attenuated FFT theta peak power. (B) Time course of FFT theta peak frequency in blocks of 5 min. Drug treatment did not evoke any statistically significant effect on frequency. Data are mean ± S.E.M. Significant difference (p < 0.05): * ‘L-733,060 MS’ group and ‘SOM MS’ group vs. the corresponding ‘Vehicle MS/LS’ groups; & ‘L-733,060 MS’ group vs. ‘L-733,060 LS’ group; two-way RM ANOVA followed by Bonferroni post-hoc test.

However, pre-treatment with L-733,060 did not affect FFT theta peak frequency (Figure 7B, left; Treatment, F_2, 75_ = 5.69, p > 0.5; two-way RM ANOVA).

#### L-733,060 did not affect exploratory theta recorded prior to formalin test

While pre-treatment with L-733,060 attenuated the power of formalin-induced theta, the drug did not significantly affect power of theta induced on exploration by the animal in test environment prior to the formalin test. The power of exploratory theta before vs. after microinjection was similar in both vehicle (0.10 ± 0.02 mV^2^ vs. 0.10 ± 0.02 mV^2^ (n=5), p > 0.7; two-tailed unpaired t-test) and L-733,060 microinjected animals (MS: 0.09 ± 0.02 mV^2^ vs. 0.08 ± 0.02 mV^2^ (n=8), p > 0.7; LS: 0.11 ± 0.02 mV^2^ vs. 0.10 ± 0.02 mV^2^ (n=10), p > 0.7; two-tailed unpaired t-test). The number of animals in each group analysed after microinjection differs from that after formalin injection into the right hind paw as the EEG recordings of a few animals following microinjection were not recorded due to technical issues.

#### Effect of microinjection of SST on formalin-induced theta activation

Two-way RM ANOVA followed by Bonferroni post-hoc test revealed that intraseptal microinjection of 3.38/6.76 μg/μl SST evoked a significant effect on theta duration (Treatment, F_2, 78_ = 7.25, p < 0.004; data not shown), normalized FFT theta peak power (Figure 7A, right; Treatment, F_2, 78_ = 4.81, p < 0.02) but not FFT theta peak frequency (Figure 7B, right; Treatment, F_2, 78_ = 2.26, p > 0.1). The change was observed in the 3.38/6.76 μg/μl SST MS group, but not the 3.38/6.76 μg/μl SST LS group, and was marked by a parallel shift in normalized power with a significant reduction of power in Phase 1. Indeed, the average of normalized power of theta in the four phases was significantly lower (Groups, F_2, 26_ = 4.81, p < 0.02; one-way ANOVA followed by Newman-Keuls post-hoc test) in the 3.38/6.76 μg/μl SST MS group (0.24 ± 0.04, n = 10) vs. Vehicle MS/LS group (0.44 ± 0.04, n = 9) and 3.38/6.76 μg/μl SST LS group (0.35 ± 0.05, n= 10).

## Discussion

The present study has led to several novel findings. One, peptide neurotransmission at NK1R, presumably involving SP in MS, modulates nociceptive responses in septo-hippocampus. Thus, on one hand, microinjection of NK1R agonist, SP, into MS evoked a robust suppression of the CA1 PS that was attenuated by intraseptal microinjection of NK1R antagonist, L-733,060. On the other hand, intraseptal L-733,060 attenuated formalin-induced suppression of CA1 PS and duration of theta activity in anaesthetized animal while attenuating the power of formalin-induced theta in behaving animal. The CA1 PS reflects the excitability of CA1 pyramidal cells to intrinsic afferent input from field CA3 while theta duration and power reflect partly the duration and population size of neurons synchronized into rhythmic theta oscillation. Notably, theta activation, theta-rhythmic excitation of CA1 pyramidal cells and suppression of CA1 PS are parts of spectrum of neural responses involved in encoding of information to salient stimulus, including formalin injection (Khanna, 1997; Lovett-Barron *et al*., 2014; Zheng & Khanna, 2001). Interestingly, the septal NKR may play a relatively selective role in mediation of septo-hippocampal network responses to salient stimuli. Thus, while intraseptal L-733,060 antagonized formalin-evoked responses, the drug failed to antagonize intraseptal carbachol-induced theta activation and suppression of CA1 PS suggesting that NK1R mechanisms are not pivotal across all forms of septo-hippocampal network activation.

The theta activity in anaesthetized rat was recorded from the pyramidal cell layer that reflects an integration of theta-rhythmic voltage changes across the cell body and proximal dendrites of pyramidal cells that are driven partly by septal cholinergic neurons (Bland, 1986; Buzsáki, 2002; Freund & Buzsáki, 1996; Leung, 1984). Since the NKRs are located almost exclusively on cholinergic neurons, the findings suggest that microinjection of L-733,060 antagonized peptidergic mediated activation of medial septal cholinergic neurons leading to a loss of theta wave activity in the hippocampal pyramidal cell region. This is also consistent with observation that the antagonist strongly attenuated formalin-induced suppression of CA1 PS, which is also mediated by cholinergic neurons of the MS (Zheng & Khanna, 2001).

Interestingly, the power and frequency of residual theta in presence of intraseptal L-733,060 were unchanged from the control suggesting a NK1R-insensitive septal mechanism equally modulates formalin-induced theta. Indeed, in behaving animal, separate microinjection of L-733,060 and SST into MS each attenuated power of formalin-induced theta wave activity recorded from dendritic regions of CA1 pyramidal cells. The power of dendritic theta in behaving animal reflects an integration of voltage changes across somatic-dendritic dipoles (Bland, 1986; Buzsáki, 2002; Freund & Buzsáki, 1996; Leung, 1984). While L-733,060 likely affected power in awake animal through modulation of the somatic component (see above), the cellular mechanism of SST effect on power is unclear, although the effect of the drug is expected to be through septal GABAergic neurons since the SSTR are localized almost exclusively on these neurons in MS (Bassant *et al*., 2005).

Two, parallel peptidergic mechanisms in MS mediate nociceptive behaviours and formalin-induced theta activation. In context of nociception, microinjection of L-733,060 into MS selectively attenuated flinching while sparing nociceptive licking in awake rat. On the other hand, microinjection of SST into MS attenuated nociceptive licking while sparing flinching. Functionally, the two drugs used at the selected doses evoked a similar level of decrease in the power of formalin-induced theta in behaving animals suggesting that they were equally effective in modulating neural activity in MS though likely through different neural mechanisms. Indeed, microinjection of L-733,060 alone into MS evoked only a ~50% decrease in duration of theta wave activity in anaesthetized rat. Complementing the neural and behavioural effects of L-733,060- and SST-sensitive mechanisms, broad-based MS inactivation with microinjection of the GABA mimetic, muscimol, that evoked a sustained loss of theta in anaesthetized animal attenuates both licking and flinching induced on injection of formalin (Lee *et al*., 2011).

The behavioural and electrophysiological effects with intraseptal SST and L-733,060, especially at peaks of the formalin test, were observed with no change in ambulation. Speed of ambulation was also unaltered on microinjection. This dissociation is notable since theta activation is also strongly associated with increased ambulation. The dissociation suggests that antinociception evoked on intraseptal microinjection of the two drugs was not secondary to ambulatory changes or sedation. Furthermore, partial changes in nociceptive behaviours *per se* do not suffice to affect theta processing, especially in context of L-733,060. Thus, microinjection of L-733,060 into both LS and MS evoked a similar decrease in flinching in the formalin test, but only the microinjection into MS attenuated theta activation. The preceding suggests that the effect of the antagonist on formalin theta reflects a direct effect of the antagonist on septo-hippocampal theta neural processing in the MS. Furthermore, the localized nature of the electrophysiological effect of L-733,060 on formalin-induced theta activation indicates that the functional spread of drug is circumscribed, thus suggesting that the behavioural effect of microinjection of L-733,060 is due to local neural elements in the region of microinjection, whether MS or LS.

Three, the localized nature of functional effects with intraseptal L-733,060 suggests that the peptide neurotransmission at NK1R is a common modulator of acute affective-motivational behaviours along the medio-lateral axis of the septum comprising the MS and LS. In this context, microinjection of L-733,060 into both the MS and LS attenuated animal ambulation in novel open field in a time-dependent fashion besides decreasing nociceptive flinching in the formalin model of persistent inflammatory pain. The drop in both behaviours was sharp.

The inhibition of locomotion in open field by intraseptal L-733,060 was most marked in the declining phase of ambulation while sparing ambulatory behaviour in the first minutes of exposure. In contrast, microinjection of SST into MS, but not LS increased ambulation during period of relatively low ambulation in the resting phase of the open field and the interphase of the formalin test. Collectively, the findings suggest that while septal NK1R mediated transmission facilitate exploratory behaviour in open field, the SST-sensitive neural mechanism in MS facilitate inactivity during habituation in open field and at interphase in the formalin test.

Consistent with a shared role of NKR in affect-motivation along the medio-lateral axis of the septum, it is notable that SP is released or its tissue levels increases in LS and MS under stressful conditions (Ebner *et al*., 2007; Siegel *et al*., 1984). The mechanism by which LS and MS may be integrated is unclear. A common peptidergic input and/or intraseptal peptidergic connections may play a role. LS presents a rich tapestry of SP afferent fibres and neurons (Risold & Swanson, 1997a, 1997b; Szeidemann *et al*., 1995). Further, neurons in dorsal and intermediate regions of LS that contain SP positive neurons also send afferent to MS region (Risold & Swanson, 1997a, 1997b; Szeidemann *et al*., 1995).

Overall, the current findings suggest that non-overlapping peptidergic NK1R and SST-sensitive neural mechanisms in septum collectively modulate (a) the processing of the noxious stimulus in septo-hippocampus and nociceptive behaviours, and (b) the sensorimotor behaviour on exposure to novel environment. Seen in juxtaposition with the role of NK1R in response to salient events in the spinal cord and ventromedial medulla (Hamity et al., 2010; Mason, 2005; Trafton et al., 2001), the current findings highlights that NK1R play a role along the of neural axis in modulating neural responses and behaviour to salient events.

The co-localization in MS of both behavioural and electrophysiological effects of L-733,060 suggests that at least in MS the cholinergic neurons comprise a shared pool of neurons for NK1R-mediated effects on formalin-induced flinching and theta activation. Indeed, consistent with the role of NK1R in salient events, the septal cholinergic neurons are implicated in registration of novelty in cingulate cortex, nociception and hippocampal theta activation (Ariffin et al., 2013; Jiang *et al*., 2018a; Khanna, 1997; Khanna & Sinclair, 1992; Zheng & Khanna, 2001). More broadly, the forebrain cholinergic neurons and the cholinergic transmission therein are implicated in both nociception and antinociception (Koga et al., 2017; Naser & Kuner, 2018; Vierck *et al*., 2016).

As regards to SST-sensitive circuit mechanisms, it is notable that microinjection of the GABA_B_ receptor antagonist, bicuculline, into MS also evoked a very robust increase in formalin-induced ambulation, including during the interphase, and a marked loss of formalin-induced theta activity (Ang *et al*., 2015). This suggests that modulation of locomotion and theta activation by MS involves in part an intraseptal release of GABA. From the perspective of a circuit mechanisms, SST may affect interphase and theta activation in part by inhibiting SSTR expressing PV^+^ GABAergic projection neurons and by intraseptal disinhibition, by inhibiting SSTR expressing local GABAergic neurons, which mimics the effect of intraseptal bicuculline. This may be basis of effect of SST in open field as well. Indeed, the PV^+^ projection neurons are implicated in decrement of locomotion and theta generation (Bender *et al*., 2015). However, of the two mechanisms, disinhibition is unlikely to mediate the effect of SST on licking since intraseptal bicuculline does not effect this behaviour (Ang *et al*., 2015). This raises a possibility that PV^+^ GABAergic projection neurons partly mediate nociceptive behaviours.

## Acknowledgements

This work was supported by research grants awarded by the National Medical Research Council, Singapore and by an award from ‘National University of Singapore, University Strategic Research (DPRT/944/09/14) and NUS Yong Loo Lin School of Medicine Aspiration Fund (R-185-000-271-720)’ to Sanjay Khanna.

---

**Figure. S1.**
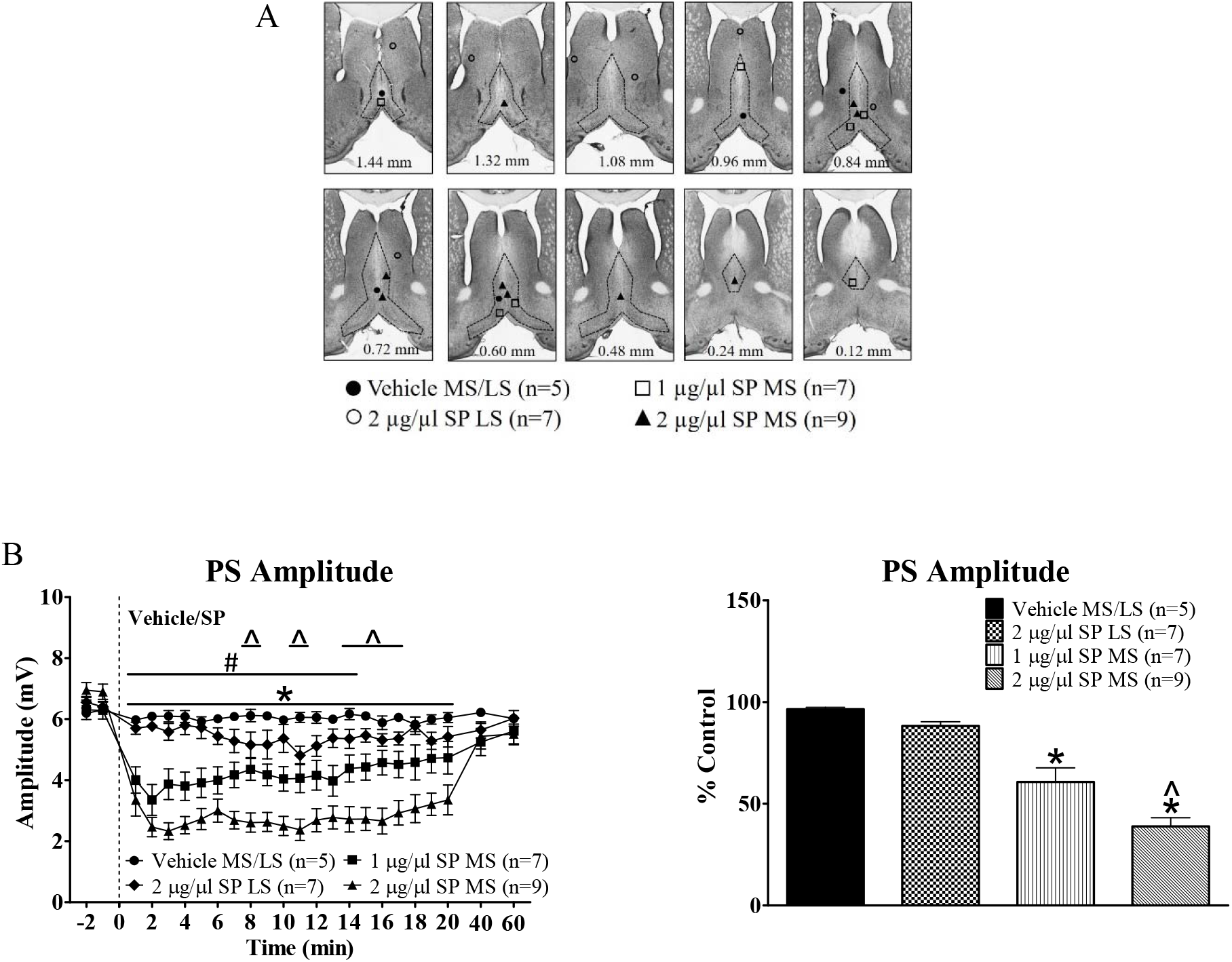
Microinjection of Substance P (SP) into the medial septum (MS) induced suppression of CA1 population spike (PS) in anaesthetised animals. (A) Composite of microinjection sites in the septal region. Microinjection was performed using a stainless steel cannula. The following drugs were microinjected in separate experiments into either MS or lateral septum (LS): Vehicle, SP (1 or 2 μg/μl, 0.5 μl). (B, left) Time course of effect of SP on the amplitude of CA1 PS. The PS was averaged in blocks of 1 min. Vehicle or SP was microinjected at time 0 min as indicated by the dashed vertical line. A given concentration of SP (or vehicle) was microinjected thrice into MS or LS with a gap of at least 1 hr between microinjections. Since the effect of repeat injections at a given dose and at the selected site were comparable, an average response for that dose and site was built by averaging the time course for the three microinjections in a given experiment and then for the entire group. (B, right) Histogram illustrating the average PS in first five minutes after microinjection. The amplitudes are expressed as percentage of the control amplitude before microinjection. Note that microinjection of SP into MS, but not LS induced a robust and consistent decrease of CA1 PS amplitude with the higher concentration of 2 μg/μl evoking a stronger suppression. Data are mean ± S.E.M. Significant difference (p < 0.05): Time course graph: * ‘2 μg/μl SP MS’ group vs. ‘Vehicle MS/LS’ group; # ‘1 μg/μl SP MS’ group vs. ‘Vehicle MS/LS’ group; ^ ‘2 μg/μl SP MS’ group vs. ‘1 μg/μl SP MS’ group; two-way RM ANOVA followed by Bonferroni post-hoc test. Significant difference (p < 0.05): Histogram: * vs. ‘Vehicle MS/LS’; ^ vs. 2 μg/μl SP LS; Kruskal-Wallis test followed by Dunn’s post-hoc test.

**Figure. S2.**
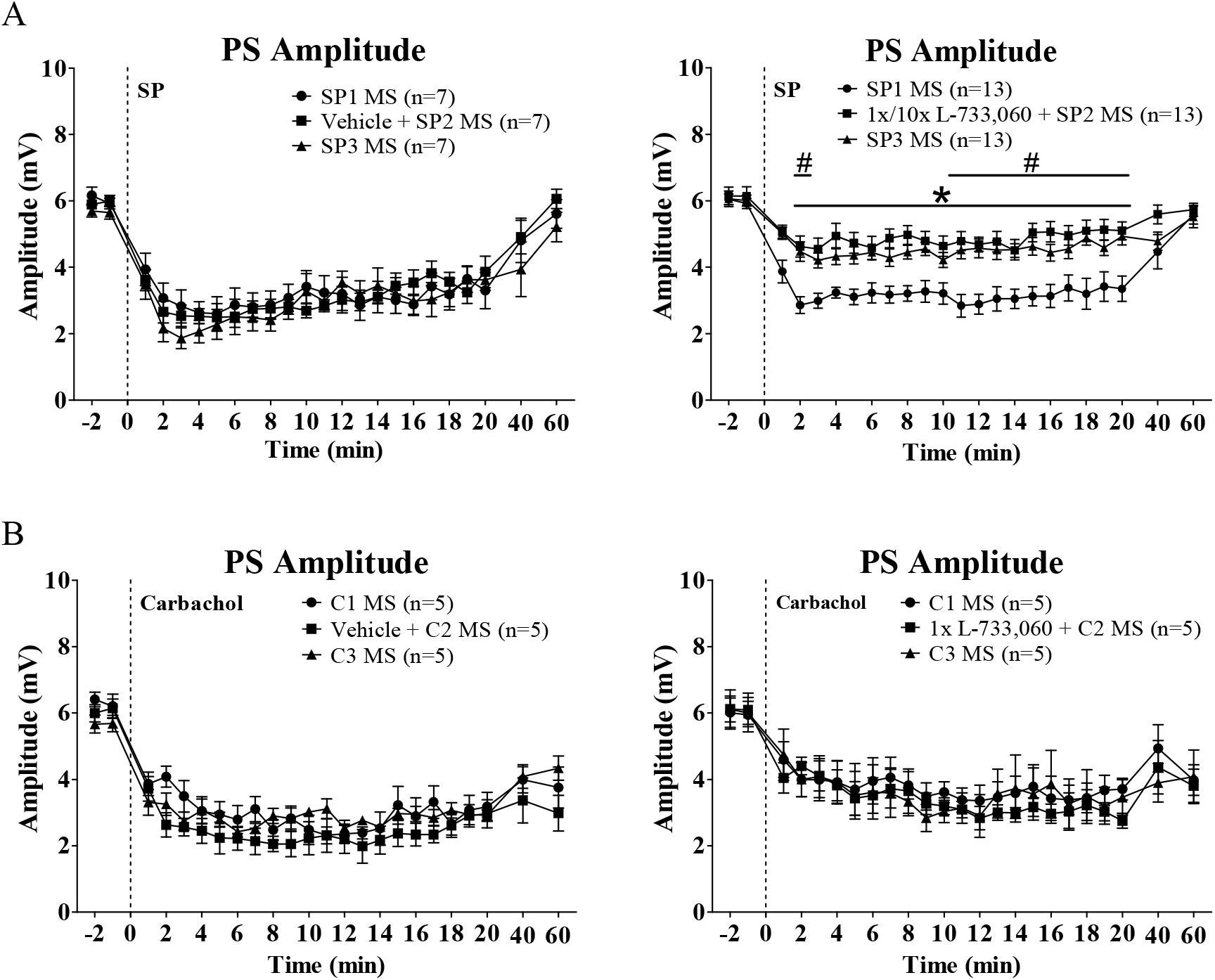
Microinjection of neurokinin 1 receptor (NK1R) antagonist, L-733,060, into the medial septum (MS) attenuated Substance P (SP)-but not carbachol-induced suppression of CA1 population spike (PS) in anaesthetized animals. Substance P (SP; 2 μg/μl, 0.5 μl) or carbachol (0.0156 μg/μl, 0.5 μl) was microinjected three times (SP1, SP2 and SP3; C1, C2 and C3) at least 1hr apart during the course of the experiment whereas vehicle or neurokinin1 receptor (NK1R) antagonist, L-733,060 (1X, 0.0176 μg/μl, 0.5 μl) was microinjected 15 min before SP2 or C2. The graphs are built as explained in Fig. S1. (A) Time course of change in PS amplitude on microinjection of SP. Microinjection of L-733,060 into MS (right) but not vehicle (left) attenuated SP-induced suppression of CA1 PS. (B) Time course of change in PS amplitude on microinjection of carbachol. Microinjection of neither vehicle (left) nor L-733,060 (right) attenuated carbachol-evoked suppression of CA1 PS. Data are mean ± S.E.M. Significant difference (p < 0.05): * ‘1x/10x L-733,060 + SP2 MS’ group vs. ‘Vehicle MS/LS’ group; # ‘SP3 MS’ group vs. ‘Vehicle MS/LS’ group; two-way RM ANOVA followed by Bonferroni post-hoc test.

